# A graph-based pangenome reveals the genetic basis of climate-resilient and horticultural traits in pear

**DOI:** 10.64898/2026.05.08.723691

**Authors:** Yuhao Gao, Wendi Wang, Yijie Liu, Jiahao Wu, Lu Wang, Jia Wei, Meisong Dai, Chunyan Wei, Luming Tian, Cuicui Jiang, Jun Su, Huabai Xue, Haiquan Liu, Junbei Ni, Shuang Jiang, Danying Cai, Xiaoyan Zheng, Dong Zhang, Songling Bai

## Abstract

Climate change poses an increasing threat to the cultivation of deciduous fruit trees, placing greater demands on modern pear breeding. Using pear germplasm adapted to diverse environments, we assembled 11 chromosome-level genomes. In combination with 13 publicly accessible pear genomes, we analyzed presence-absence variations (PAVs) and constructed a graph-based pangenome for pear. By performing a PAV-eQTL analysis of the fruit of 123 pear accessions, we identified PAVs significantly associated with expression levels of genes that may be involved in regulating agronomic traits. Population analysis of 268 pear accessions revealed two stop-gained variants in *DAM1* of independent origin, which may function in advancing the blooming date and reducing the chilling requirement. We detected complex PAVs at the *NOR1* locus, including two copy-number variations and one deletion. These PAVs contributed to the rapid diversification of the *NOR1* locus and the fruit development period through regulating *ARF5* and other ripening-related genes. We revealed the selection history of the *NOR1* locus and developed novel pear individuals that accumulated alleles for low chilling requirement, early blooming date, and short fruit development period. The results provide valuable resources for pear genomics research and offer a guideline for breeding modern pears with climate resilience.

## Introduction

Global climate change is fundamentally disrupting the phenological synchronization of temperate perennial crops, threatening the sustainability of fruit production systems worldwide. Deciduous fruit trees, such as pear (*Pyrus* L.), rely on a delicate balance of environmental cues–specifically winter chill accumulation and spring heat–to regulate the dormancy cycle and flowering period. Failure to satisfy the chilling requirement (CR) attributable to warmer winters leads to erratic blooming and unstable yield^1^, necessitating the urgent development of climate-resilient cultivars. Concurrently, modern horticulture demands cultivars with an optimized fruit development period (FDP) and ripening characteristics compatible with supply chain logistics and consumer preferences. However, the genetic architecture underlying these complex phenological and quality traits remains obscure owing to the high heterozygosity and long generation times characteristic of woody perennials.

Single-nucleotide polymorphisms (SNPs) have traditionally dominated the dissection of complex traits. Recent studies of major crops have revealed that structural variations (SVs), which includes presence/absence variations (PAVs) and copy-number variations (CNVs, a specific type of PAV), are pivotal drivers of rapid evolutionary adaptation and domestication^2^. In species ranging from tomato^3^ to apple^4^, SVs have been shown to modulate gene expression through dosage effects, often explaining phenotypic variance that SNPs fail to capture. However, the complexity of SVs complicates their characterization. In recent years, graph-based pangenomes, which integrate SV information that effectively bridges this gap, have resolved the genetic basis for critical traits in plants^5,6,7^. However, in the genus *Pyrus*, a comprehensive analysis of SVs and a graph-based pangenome are lacking, hampering the exploration of causal alleles for agronomic traits.

Here, we present a graph-based pangenome for pear, constructed from 24 chromosome-level assemblies, including 11 assemblies from nine newly sequenced accessions representing diverse ecotypes. By integrating multi-omics data with population-scale phenotyping, we dissect the genomic footprint of natural adaptation and artificial selection. We identify two distinct classes of genomic variants that drive evolution in crucial horticultural traits: (1) convergent loss-of-function mutations in the *Dormancy-Associated MADS-box 1* (*DAM1*) locus, whereby independent stop-gain variants act as evolutionary switches to lower the CR in warm-climate-adapted landraces; and (2) gene dosage expansion of the *Non-Ripening 1* (*NOR1*) transcription factor, where stepwise increase in copy number is causally linked to accelerated fruit ripening and shortened FDP. These findings provide insight into the genetic basis of critical phenological traits and present a genomic design framework for breeding early-maturing, low-CR pear cultivars that are resilient to a warming climate.

## Results

### Eleven chromosome-level genome assemblies provide a high-quality pear genomic foundation

Nine accessions from four *Pyrus* specie were selected for genome sequencing (Supplementary Table 1). ‘Yali’ (YL), ‘Golden Nijisseiki’ (J20), ‘Qinghua’ (QH) and ‘Wonhwang’ (YH) are cultivars of *P*. *pyrifolia* (sand pear); ‘Bartlett’ (BL) is a cultivar of *P*. *communis* (European pear) cultivar; ‘Nanhong’ (NH) is a *P*. *ussurensis* cultivar; ‘Hongchuanli’ (HCL) represents a *P*. *pashia* accession; ‘Meili’ (ML) is a *P*. *pyrifolia* landrace that needs to ripen after harvest; and ‘Yunnanshali’ (YNSL) is likely a hybrid of *P*. *pyrifolia* and *P*. *pashia* (Supplementary Note 1 and Supplementary Fig. 1).

We generated 25.98–81.78 Gb of Oxford Nanopore Technologies (ONT) long reads for seven accessions (YL, ML, J20, NH, YNSL, HCL and BL) and 29.7–38.0 Gb of PacBio HiFi reads for QH and YH (Supplementary Table 1). For each accession, we additionally generated 35.4–61.1 Gb of chromosome conformation capture (Hi-C) reads (Supplementary Table 1). We assembled haploid consensus assemblies for accessions with ONT reads and haplotype-resolved assemblies for QH and YH (designated QH_H1, QH_H2, YH_H1 and YH_H2). All 11 assemblies were anchored to chromosome-level genomes using the Hi-C reads (Supplementary Fig. 2). The contig N50 of the newly generated genome assemblies ranged from 1.96–28.8 Mb (Supplementary Table 2). BUSCO evaluation revealed 97.4%–99.0% complete BUSCOs in the newly assembled genomes (Supplementary Table 2). The long terminal repeat (LTR) assembly index (LAI)^8^ for every genome exceeded 18 (Supplementary Table 2). These results indicated that the newly assembled genomes were of high quality.

Approximately 50.95%–52.84% of the genomes were annotated as repeat sequences (Supplementary Table 3). Retroelements and DNA transposons comprised the two major types of repeat sequences and the total length of retroelements exceeded that of DNA transposons (Supplementary Fig. 3 and Supplementary Table 3). In the 11 genomes, we identified 3,069–4,250 intact LTR-RTs (Supplementary Fig. 4a) that were inserted during the last 2 million years (Supplementary Fig. 4b). We annotated 40,891–46,857 gene models in the 11 genomes with BUSCO values of 93.8%–98.0% (Supplementary Table 2).

### A graph-based pangenome captures extensive diversification driven by PAVs

To further capture the diversity of the pear genomes, we integrated 13 published pear genomes (Supplementary Table 4) with the newly assembled 11 genomes, representing five *Pyrus* species. In total, 48,322 gene clusters were identified in the 24 genomes and were designated as pan–gene clusters. Of these gene clusters, 24.3% were designated core clusters shared by all genomes, 73% were shared by 2–23 genomes and were considered to be shell clusters, and 2.7% were present in one genome only and were designated private clusters (Fig. 1a). We further categorized the genes in each genome based on the gene cluster analysis. Approximately 40%–54% of the genes in each genome were core genes, whereas shell genes comprised 43%–56% of the total genes (Fig. 1b). We compared the characteristics of the three types of gene using YL assembly as an example. Approximately 87.5% of the core genes were successfully annotated from the Pfam database, exceeding the percentages for the shell and private genes (Fig. 1c). The ratio of nonsynonymous substitutions to synonymous substitutions (*K_a_/K_s_*) for the core genes was much lower than that for the shell genes (Fig. 1d). In addition, the coding sequence (CDS) length and expression levels of the core genes were higher than those of the shell and private genes (Supplementary Fig. 5). These results indicated that the core genes were considerably more functionally conserved than the other gene types.

**Fig. 1.**
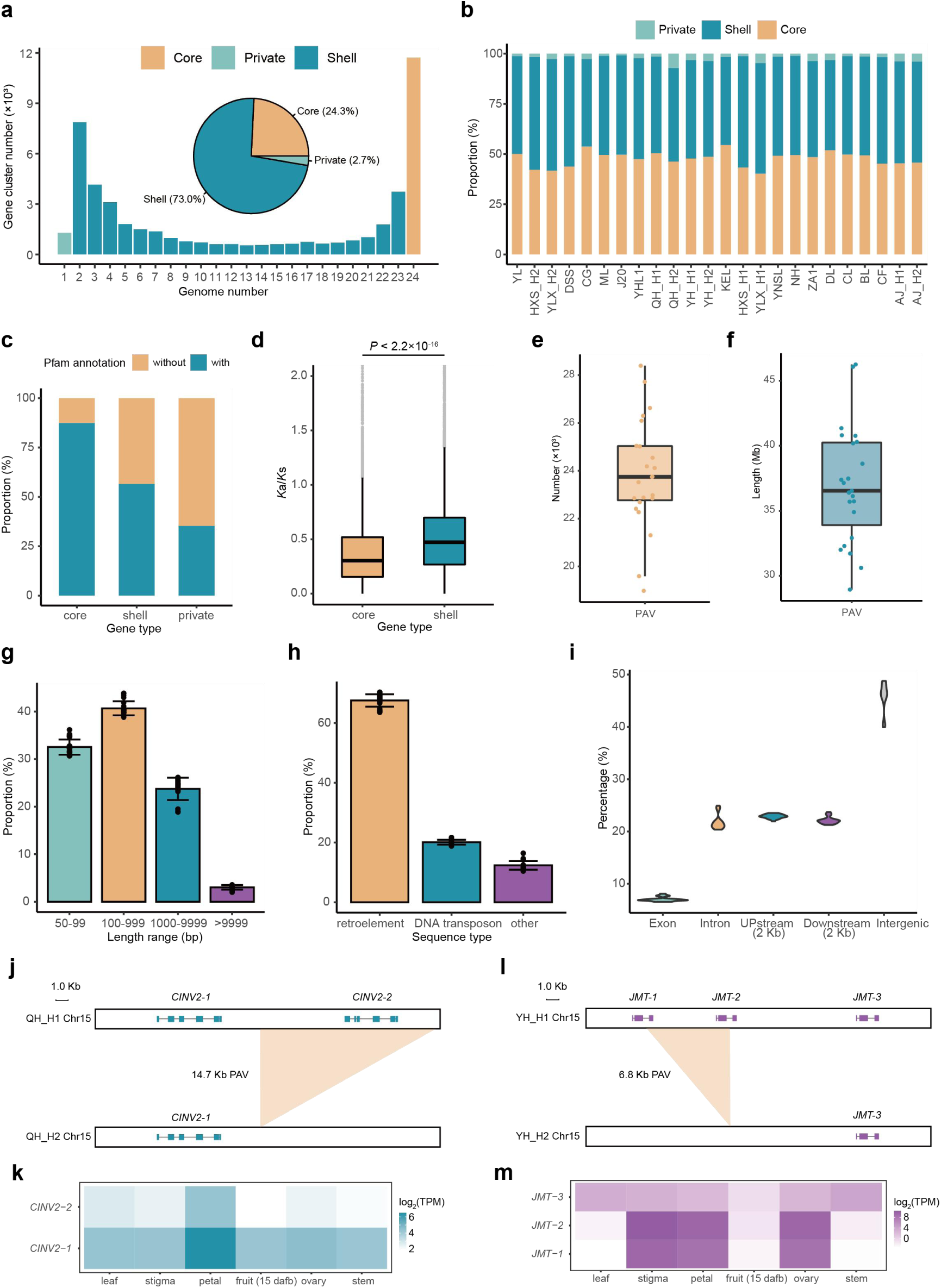
Pangenome analyses based on 24 pear genomes. **a** The pie chart shows the composition of core, shell, and private gene cluster among all clusters. The bar chart indicates the number of gene clusters across 24 genomes at different frequencies. **b** The proportion of core, shell and private genes in each genome. **c** The proportion of core, shell and private genes with Pfam annotation. **d** *Ka*/*Ks* values of core genes were lower than those of shell genes (two-sided *t*-test, zoom in to 0–2 of y axis). **e** Total number of PAVs in each genome. **f** Total base length of PAVs in each genome. **g** Base length distribution of PAVs in each genome. **h** Repeat sequence annotation of PAVs in each genome. **i** A violin plot showing the percentage of PAVs overlapped with different genomic features. **j** A PAV leading to the loss of *CINV2-2*. Colored rectangles represent exons. **k** Expression levels of *CINV2-1* and *CINV2-2* in different organs. **l** A PAV leading to the loss of coding regions of *JMT-1* and *JMT-2*. Colored rectangles represent CDSs. **m** Expression levels of *JMT-1*, *JMT-2* and *JMT-3* in different organs. In boxplots, the 25% and 75% quartiles are shown as the lower and upper edges of the boxes, respectively, and central lines denote the median. The whiskers extend to 1.5× the interquartile range. Data in **g** and **h** are given as mean ± s.d.

To analyze the PAVs in the pear genome, we aligned 23 pear assemblies to the YL assembly. We detected 18,985–28,389 PAVs (Fig. 1e) in each genome with the total bases ranging from 28.9–46.3Mb (Fig. 1f). Most PAVs were shorter than 10 Kb (Fig. 1g) and, on average, 67.6% and 20.1% of the PAV sequences were annotated as retroelements and DNA transposons (Fig. 1h), indicating transposable elements are a major component driving the modification of the pear genome. All the PAVs were merged into 178,687 non-redundant records and a graph-based pangenome was constructed.

Approximately 7.1% and 22.9% of PAVs were located in exons and 2,000 bp promoter regions upstream of genes, respectively (Fig. 1i), indicating that they potentially affect gene structure and expression. For example, a 14,271 bp PAV resulted in the loss of one copy of *CINV2* (encoding a cytosolic invertase) in the QH_H2 haplotype (Fig. 1j and Supplementary Fig. 6). Because the two *CINV2* copies (*CINV2-1* and *CINV2-2*) were expressed in several tissues (Fig. 1k), the loss of one copy may affect carbon partitioning^9^ in plants. Two *JMTs* genes (*JMT-1* and *JMT-2*, encoding jasmonic acid carboxyl methyltransferases) were absent in YH_H2 but present in YH_H1, which was caused by a 6,773 bp PAV overlapping with the CDS regions (Fig. 1l and Supplementary Fig. 7). JMTs catalyzed the conversion of jasmonic acid to the methyl jasmonate, thus affecting the content of the latter in plants^10^. RNA-Seq revealed that the *JMTs* were expressed in reproductive organs (Fig. 1m), which may affect seed development^11^ through regulating dosage of methyl jasmonate.

### Gene expression divergence and associated PAVs

The predominant cultivated pear species, *P*. *communis* and *P*. *pyrifolia* exhibit notable differences in fruit traits. By analyzing RNA-sequencing (RNA-Seq) data from the fruit of 38 *P*. *communis* and 50 *P*. *pyrifolia* cultivars, we detected 2,929 genes that were significantly differentially expressed between the two species (Fig. 2a). For example, *PG1*, a critical gene influences fruit firmness by catalyzing cell wall degradation^12^, was expressed at significantly higher levels in *P*. *communis* than in *P*. *pyrifolia* (Fig. 2b), which may contribute to the softer texture of *P*. *communis* fruit. Genes potentially associated with fruit sugar and acid accumulation^13,14^ (such as *SWEET15*, *ERDL6*, *SUT4*, *NADP-ME1* and *Ma1*) showed significant differences in expression level between *P*. *communis* and *P*. *pyrifolia* (Fig. 2c, d and Supplementary Fig. 8), suggesting that the genetic basis of the regulation of fruit quality traits may differ in the two species. Notably, a previously identified PAV leading to the loss of *Ma1*^15^ may explain the lower *Ma1* expression level in *P*. *pyrifolia*. However, when repeating the analysis using *P*. *pyrifolia* accessions lacking this PAV, the expression differences remained significant (Supplementary Fig. 9a). The promoter sequence of *Ma1* was strongly divergent between *P*. *communis* and *P*. *pyrifolia* which was mainly caused by a PAV (Supplementary Fig. 9b, c). Dual-luciferase reporter assays indicated that the activity of the *Ma1* promoter in *P*. *communis* is significantly higher than that in *P*. *pyrifolia* (Fig. 2e), indicating that differences in promoter activity may be one reason for the differential expression of *Ma1*.

**Fig. 2.**
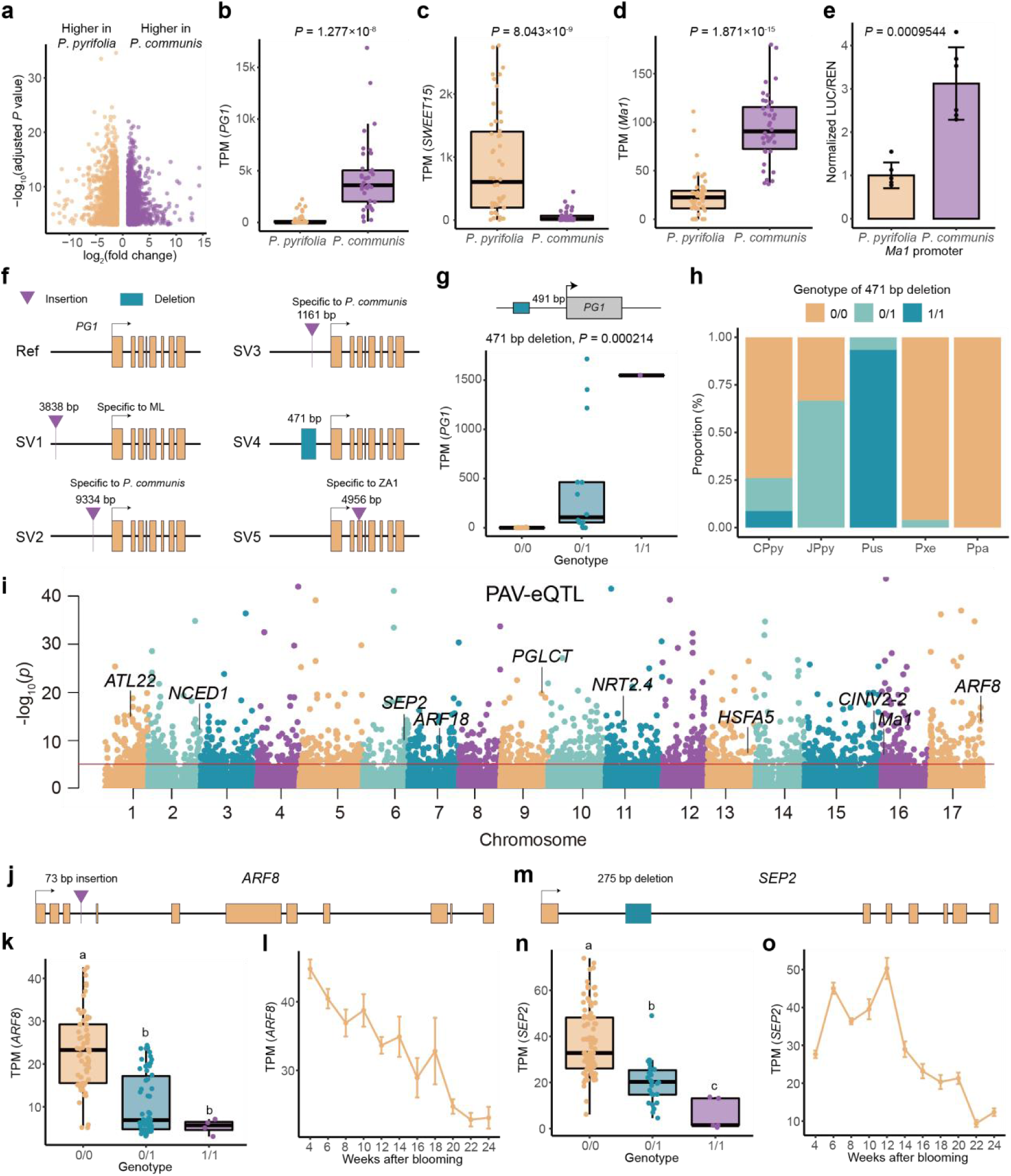
Gene expression divergence and associated PAVs. **a** Differentially expressed genes between *P*. *communis* and *P*. *pyrifolia* fruit. **b-d** The expression levels of *PG1* (**b**), *SWEET15* (**c**) and *Ma1* (**d**) differed between *P*. *communis* and *P*. *pyrifolia* fruit. **e** The *Ma1* promoter from *P*. *communis* exhibits higher activity than the one from *P*. *pyrifolia*. *P* values in **b**, **c**, **d** and **e** were determined by two-sided *t*-test. **f** Five SVs at the *PG1* locus. **g** The genotype of a 471 bp deletion was significantly associated with the expression level of *PG1* in *P*. *pyrifolia* fruit. *P* values were determined by linear regression model. **h** Frequency of the 471 bp deletion among different pear accessions (CPpy: Chinese *P*. *pyrifolia*, JPpy: Japanese *P*. *pyrifolia*, Pus: *P*. *ussuriensis*, Pxe: *P*. *xerophila*, Ppa: *P*. *pashia*). **i** Manhattan plot of association between PAV genotype and expression level of adjacent gene in developing fruit of *P*. *pyrifolia* accessions. The red line indicates the significance cutoff (*P* = 1 × 10^-5^). **j** A 73 bp insertion was detected in an intron of *ARF8*. **k** The genotype of the 73 bp insertion was significantly associated with the expression level of *ARF8* in developing fruit of *P*. *pyrifolia* accessions. **l** *ARF8* expression during fruit development in ‘Dangshansu’ (DSS). **m** A 275 bp deletion was detected in an intron of *SEP2*. **n** The genotype of the 275 bp deletion was significantly associated with the expression level of *SEP2* in developing fruit of *P*. *pyrifolia* accessions. **o** *SEP2* expression during fruit development in DSS. In boxplots, the 25% and 75% quartiles are shown as the lower and upper edges of the boxes, respectively, and central lines denote the median. The whiskers extend to 1.5× the interquartile range. Different letters in **k** and **n** indicate statistically significant differences (*P* < 0.05, one-way ANOVA with least significant difference (LSD) test). Data in **l** and **o** are given as mean ± s.d. with three biological replicates.

To investigate the basis for the differences in *PG1* expression between *P*. *communis* and *P*. *pyrifolia*, we generated RNA-Seq data from the fruit of an interspecific hybrid (between *P*. *pyrifolia* and *P*. *communis*) and found only the *PG1* allele from *P*. *communis* was expressed (Supplementary Fig. 10). Analysis of the *PG1* promoter sequences from *P*. *pyrifolia* and *P*. *communis* revealed two insertions (9,334 bp and 1,161 bp) that were fixed in *P*. *communis* (Fig. 2f, Supplementary Fig. 11). In addition, we detected another three SVs at the *PG1* locus (Fig. 2f). A 471 bp deletion was significantly associated with *PG1* expression in the fruit of *P*. *pyrifolia* accessions (Fig. 2g). Using fruit RNA-Seq data from a cultivar heterozygous for this deletion, only the allele with the 471 bp deletion was expressed (Supplementary Fig. 12), supporting the hypothesis that allele diversification led to variation in *PG1* expression. Genotyping of different pear groups showed that this deletion has a relatively high allele frequency in the *P*. *ussuriensis* population (which produces soft fruit) (Fig. 2h), suggesting that this active *PG1* allele may have been introgressed from *P*. *ussuriensis* into *P*. *pyrifolia*. Regarding the other two SVs, a 3,838 bp insertion (specific to ML) was located in the promoter of *PG1* and a 4,956 bp insertion (specific to ZA1) was located in an exon of *PG1* (Fig. 2f, Supplementary Fig. 11). Thus both SVs may affect *PG1* transcription or translation. On the basis of these results, we propose that the polymorphism of *PG1* drives the differential *PG1* expression in which SVs may play a role, ultimately leading to heterogeneity in pear fruit firmness.

Given that SVs significantly affect gene expression^16^, we analyzed the association between PAV genotype and the expression of adjacent genes in developing fruit of 123 *P*. *pyrifolia* accessions. The pipeline identified 1,892 PAVs that were significantly associated with the expression of adjacent genes (*P* < 1 × 10^-5^) (Fig. 2i), including the two large deletions that led to the loss of *CINV2-2* (Fig. 1j) and *Ma1*^15^. A 73 bp intronic PAV was significantly associated with the expression level of *ARF8* (Fig. 2j, k), which encodes an auxin response factor. Homologs of *ARF8* function in the regulation of fruit initiation in *Arabidopsis*^17^ and likely regulate parthenocarpy in tomato^18,19^. The RNA-Seq data indicated that *ARF8* expression gradually decreased during pear fruit development (Fig. 2l). Similarly, a 275 bp intronic PAV was significantly associated with the expression level of *SEP2*, which encodes a MADS-box transcription factor (Fig. 2m, n). Cucumber *CsSEP2* functions in regulating fruit length^20^, while knock-out of a homolog of *AtSEP2* in tomato affects fruit development^21^. During pear fruit development, *SEP2* exhibited a gradually down-regulated expression pattern (Fig. 2o). Given that *ARF8* and *SEP2* have conserved functions in regulating fruit development in different species, we propose that the differential expression among different alleles of *ARF8* and *SEP2* in pear fruit may affect fruit initiation and other fruit development processes. However, further study is required to elucidate the causal variants responsible for these differences in expression patterns.

### Convergent evolution of stop-gained variations in *DAM1* confers adaptation to warm winters

The BD determines when fruit development begins. An earlier BD is beneficial for earlier fruit maturation under a suitable environment. The BD varies substantially among pear accessions (Fig. 3a). To clarify the genomic basis for the early BD trait, we reanalyzed 268 resequenced pear accessions from public data^22^ using our newly assembled genome as the reference. A genome-wide association study (GWAS) analysis revealed a significant locus for BD on Chr08 (Fig. 3b), which is located in the region of three *DAM* genes (Fig. 3c). Considering that *DAM* genes are responsible for BD and CR in rosaceous species^23^, we identified two stop-gained SNPs in the second exon of *DAM1* (Fig. 3d). These two *DAM1* alleles with stop-gained SNPs were designated *dam1-1* and *dam1-2*, respectively. We found the SNPs among *dam1-1*, *dam1-2* and *DAM1* (from Japanese sand pear) were comparable (Fig. 3e), suggesting that *dam1-1* and *dam1-2* may have independent origins. To analyze the effect of these two mutations, we genotyped 262 pear accessions and determined that the accessions harboring two *dam1* alleles (*dam1-1* or *dam1-2*) significantly showed earlier BD than the accessions with one or no *dam1* allele (Fig. 3f), indicating that *dam1* alleles may have an additive effect.

**Fig. 3.**
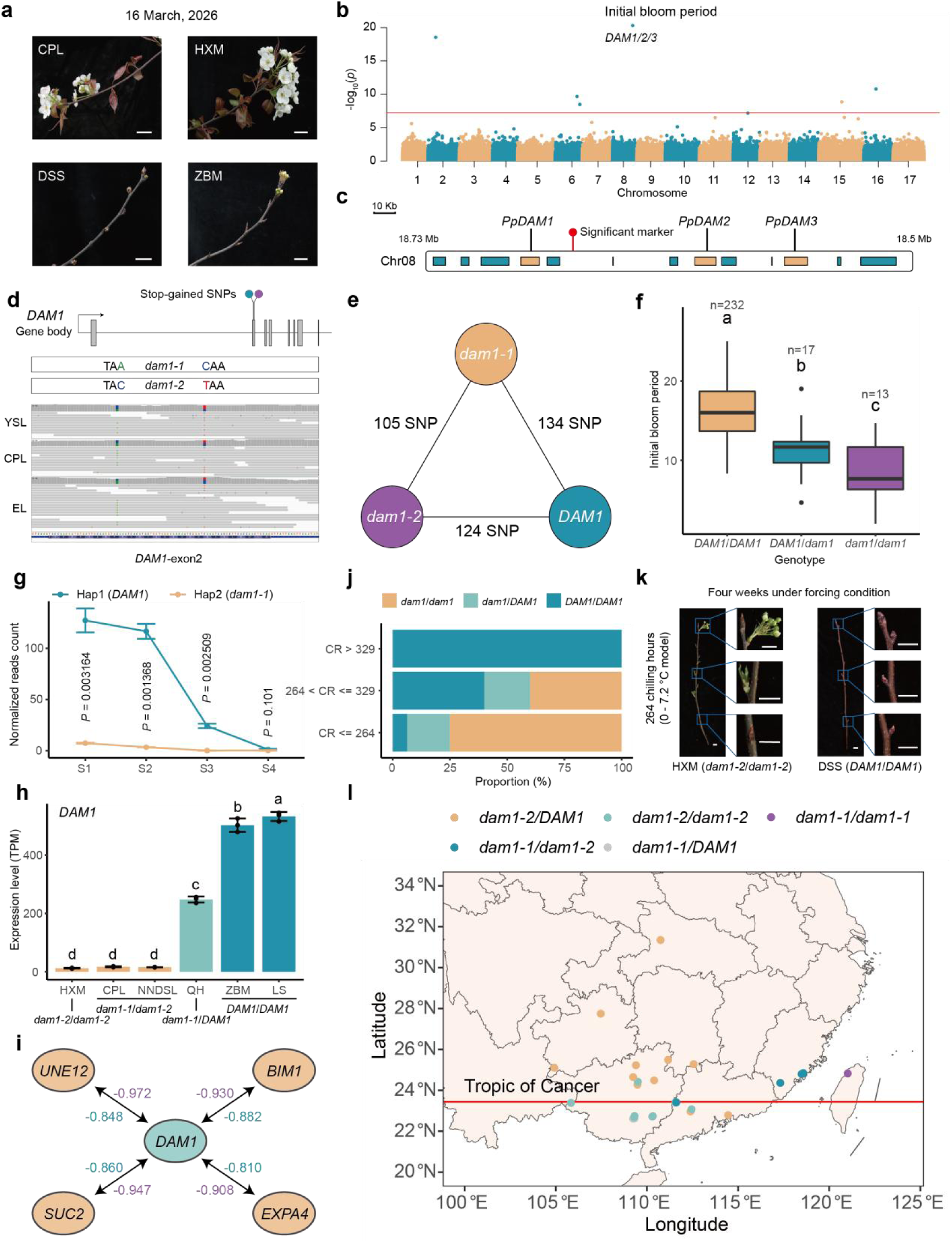
Convergent evolution of stop-gained variations in *DAM1* confers adaptation to warm winters. **a** Photographs of sand pears with different BD. Scale bar: 3.8 cm. **b** Manhattan plot for GWAS results for BD using the InDel markers and the FarmCPU model. The red line indicates the significant cutoff calculated as 0.05/*n*, where *n* is the total marker number. **c** The candidate region controlling BD contains three *DAM* genes. **d** Two independent stop-gained SNPs in *DAM1*. **e** SNP numbers among *dam1-1*, *dam1-2* and *DAM1* (from Japanese sand pear). **f** Association between the number of stop-gained *DAM1* allele (*dam1*) and BD. The 25% and 75% quartiles are shown as the lower and upper edges of the boxes, respectively, and central lines denote the median. The whiskers extend to 1.5× the interquartile range. **g** Allelic expression of *DAM1* in QH (*dam1-1*/*DAM1*) buds at four stages (S1:pre-chilling accumulation, S2: early chilling accumulation, S3: late chilling accumulation, and S4: pre-bud break). *P* values were determined by two-sided *t*-test. **h** *DAM1* expression in buds of different accessions with different *DAM1* genotypes. Sampling date: 22 November, 2025. **i** Genes with expression levels highly negatively correlated with *DAM1*. Numbers represent Pearson correlation coefficients calculated from RNA-Seq data. Same-color numbers share the same dataset. **j** Proportion of different *DAM1* genotypes among accessions with varying CR. **k** Photographs of two accessions with extremely low CR (≤ 264 units) and medium/high CR (> 329 units). Scale bar: 2 cm. **l** Geographical distribution of accessions carrying *dam1-1* or *dam1-2*. Different letters in **f** and **h** indicate statistically significant differences (*P* < 0.05, one-way ANOVA with LSD test). Data in **g** and **h** are given as mean ± s.d. with three biological replicates.

Analysis of the RNA-Seq data for QH (*dam1-1*/*DAM1*) buds collected from pre-chilling accumulation stage to pre-bud break revealed that the expression of *dam1-1* was almost undetectable (Fig. 3g). The RNA-Seq data for six cultivars differing in *DAM1* genotype showed that the expression levels of both *dam1-1* and *dam1-2* were low (Fig. 3h). The low expression of *dam1-1* and *dam1-2* may reflect the decay of nonsense mRNA, a widely observed and intensively studied phenomenon in eukaryotic cells^24^. Given that DAM1 contained an EAR motif which mediates transcriptional repression^25^, genes such as *UNE12*, *BIM1*, *SUC2*, and *EXPA4*, whose expression levels were highly negatively correlated with *DAM1* (Fig. 3i, Supplementary Fig. 13), were candidate downstream targets of DAM1.

We evaluated the CR for 42 accessions and categorized them as extremely low CR (CR ≤ 264 units), low CR (264 units < CR ≤ 329 units) or medium/high CR (CR > 329 units) cultivars (Supplementary Fig. 14). Genotyping of these accessions showed that all accessions carrying the *dam1* were classified as extremely low or low CR (Fig. 3j, k), indicating that the *dam1-1* and *dam1-2* alleles may be responsible for the low CR trait. Assessment of the native habitats of landraces revealed that landraces carrying *dam1-1* or *dam1-2* are predominantly native to areas in Guangdong, Guangxi, and Fujian provinces, which situated close to the Tropic of Cancer (Fig. 3l). The winter in these areas is warm, which is unsuitable for the growth and reproduction of pears with a high CR genotype. Thus, we hypothesize that *dam1* alleles may lead to a low CR, promoting the adaptation of pear to a warm-winter environment.

### Copy-number variations of *NOR1* contribute to a short fruit development period

The FDP is a crucial factor determining the harvest date of pear fruit. Records revealed that the FDP of improved cultivars is significantly shorter than that of landraces (Fig. 4a). The genetic distance (*F*_st_) based on PAVs (Supplementary Fig. 15) indicated that a 150 Kb region on Chr03 is highly differentiated between landraces and improved cultivars (Fig. 4b), indicating that PAVs may contribute to the variation in FDP. Reanalysis of public genomic data for two sand pear populations^22,26^ revealed the differentiated region was co-localized with the association signals for fruit harvest date and FDP (Fig. 4c and Supplementary Fig. 16). Within the differentiated region, we identified a 74,016 bp insertion whose genotype was significantly associated with FDP (Fig. 4d, e). This insertion was downstream of a NAC transcription factor and additional two intact NAC transcription factors were located within the insertion (Fig. 4d). These three NAC transcription factors were homologs of *SlNOR* in tomato (Supplementary Fig. 17), which is a critical gene promoting fruit ripening^27^, and thus we named them *NOR1*, *NOR2*, and *NOR3*, respectively. By analyzing the variation in short read depth and conducting DNA-qPCR analysis, we validated the CNV in cultivars with a high copy number for *NOR1* (Fig. 4f, g) deduced from diploid genome assemblies. *NOR1* was expressed in different reproductive organs (Supplementary Fig. 18) and was up-regulated during fruit development (Fig. 4h), indicating that *NOR1* and its copies may participate in regulating fruit ripening of pear.

**Fig. 4.**
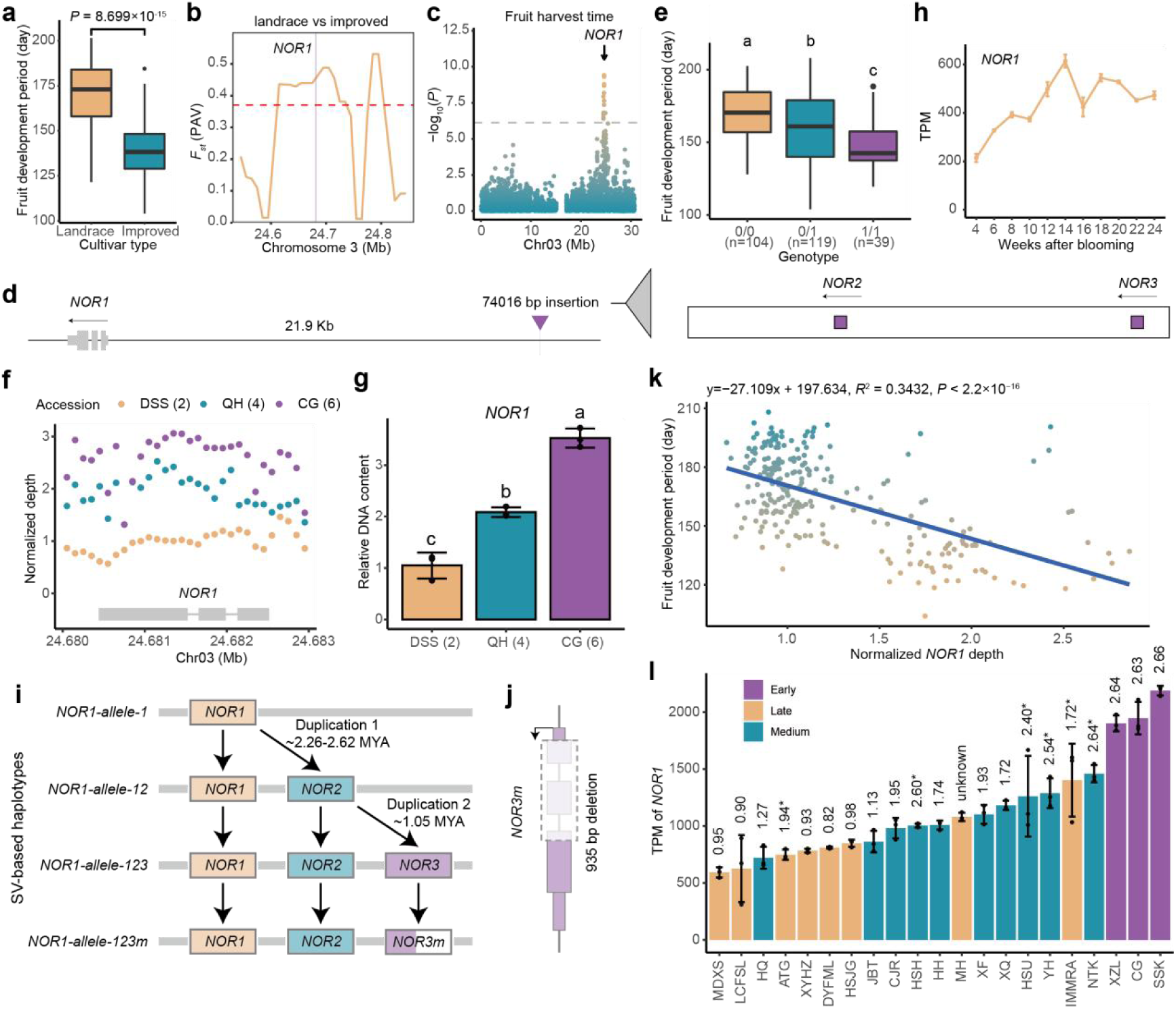
Copy-number variations of *NOR1* contribute to a short fruit development period. **a** FDP of improved cultivars (*n* = 40) is significantly shorter than that of landraces (*n* = 173). *P* value was determined by two-sided *t*-test. **b** Population divergence (*F*_st_) of landraces and improved cultivars based on PAV genotypes on Chromosome 3. The red line indicates the top 1% *F*_st_ threshold. The purple line indicates the gene body of *NOR1*. **c** Manhattan plot showing the GWAS results for fruit harvest time on Chromosome 3 using SNP markers and the mixed linear model. The arrow indicates the location of *NOR1*. The red line indicates the significant cutoff calculated as 0.05/*n*, where *n* is the total marker number. **d** A 74 Kb PAV containing two additional *NOR1* copies was identified 21.9 Kb upstream of *NOR1*. **e** The genotype of the 74 Kb PAV was significantly associated with FDP. **f** Normalized sequencing depth of different cultivars around *NOR1* gene body. Each point represents a 100 bp window. **g** DNA-qPCR validation of the additional *NOR1* copies. In **f** and **g**, the numbers in parentheses represent the known number of *NOR1* copies in diploid assemblies. **h** Expression pattern of *NOR1* during fruit development in DSS. **i** A profile showing the SV-based haplotypes of *NOR1* locus. **j** Details of the PAV on *NOR3*. **k** Association between normalized *NOR1* depth and FDP. The blue line indicates the linear regression line. The corresponding formula, *R^2^* and *P* value are showed at the top. **l** The expression levels of *NOR1* in fruit of different cultivars with different maturing dates. The numbers above the columns represent normalized *NOR1* depth. The asterisks indicate the cultivars carrying *NOR1-allele-123m*. Data in **h** and **l** are given as mean ± s.d. with three biological replicates. Data in **g** are given as mean ± s.d. with three technical replicates. Different letters in **e** and **g** indicate statistically significant differences (*P* < 0.05, one-way ANOVA with LSD test).

In the newly assembled genomes, we identified the haplotypes with one or three *NOR1* copies, and the haplotype with two *NOR1* copies was assembled using 57.8 Gb of newly sequenced PacBio HiFi reads from ‘Hongfen’. The YH_H2 haplotype had an allele with three *NOR1* copies, but the presence of a deletion at *NOR3* led to the loss of the first two exons (Fig. 4j). Totally, we identified four SV-based haplotypes, which we named *NOR1-allele-1*, *NOR1-allele-12*, *NOR1-allele-123*, and *NOR1-allele-123m* (Fig. 4i). We estimated that the first and second duplication of *NOR1* occurred approximately 2.26–2.62 million years ago (MYA) and 1.05 MYA, respectively. These duplication events greatly post-dated the divergence of *Pyrus* and *Malus*^28^, indicating their specificity to *Pyrus*.

Using the normalized depth of *NOR1* to represent the total copy number of *NOR1*, FDP was negatively associated with *NOR1* copy number at the population level (Fig. 4k). RNA-Seq data showed that early-maturing cultivars have a high *NOR1* copy number and higher *NOR1* expression in the fruit (Fig. 4l). These results suggested the CNV of *NOR1* may shorten the FDP through a dosage effect.

### Copy-number variations of *NOR1* modulate the fruit development period through dosage effects

As typical transcription factors, NOR1 and its copies may shorten fruit FDP by regulating the expression of downstream genes. To clarify the downstream genes, we conduct DNA affinity purification sequencing (DAP-Seq) of NOR1. In total, 13,517 credible peaks were enriched at the transcription start site (Fig. 5a) and the conserved binding motif for the NAC transcription factor family^29^ was enriched in the center of these peaks (Fig. 5b and Supplementary Fig. 19). Approximately 19.8% of the peaks intersected with the promoter regions (Fig. 5c), resulting in 2,342 candidate downstream genes for NOR1, among which the fruit-ripening-related genes *NADP-ME3*^30^ and *ARF5*^31^ were identified, together with a new transcription factor AGL61 (Fig. 5d). *AGL61* regulates seed development in *Arabidopsis*^32^ and may be involved in regulating fruit ripening. These target genes were up-regulated in fruit of ‘Cuiguan’ (a typical early-maturing cultivar with the genotype *NOR1-allele-123*/*NOR1-allele-123*), which were in line with the total expression level of all *NOR1* copies (Fig. 5e). Dual-luciferase assays showed that NOR1, NOR2 and NOR3 could activate the expression of *NADP-ME3*, *ARF5*, and *AGL61* (Fig. 5f), indicating the functional conservation of different *NOR1* copies in regulating specific ripening-related genes, and suggesting that a dosage effect of *NOR1* is responsible for a shortened FDP.

**Fig. 5.**
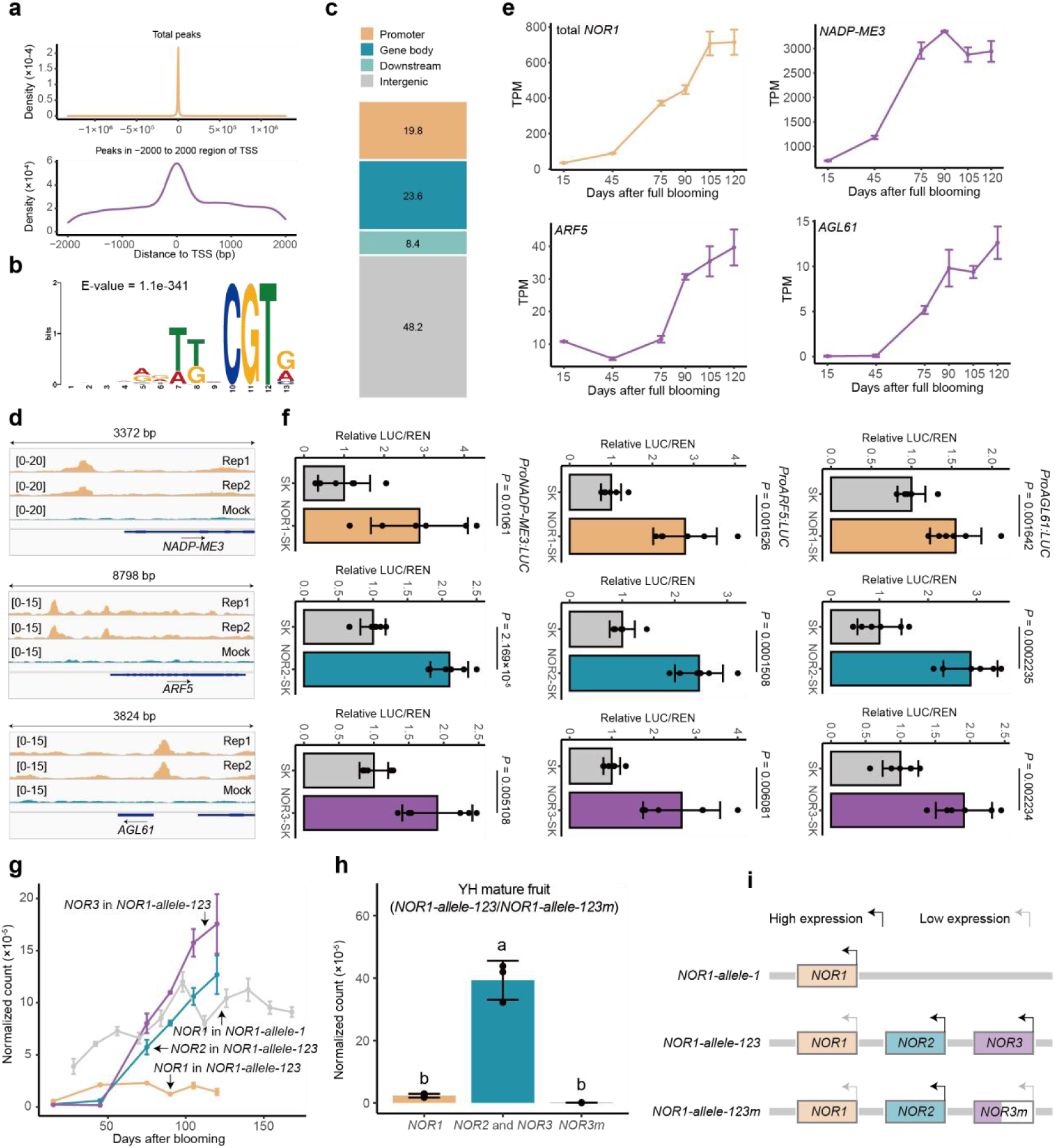
Copy-number variations of *NOR1* modulate the fruit development period through dosage effects. **a** The binding peaks identified by DAP-Seq of NOR1 are enriched around transcription start site (TSS). **b** DNA sequence motif of the NOR1-binding site. **c** Distribution of NOR1-binding peaks across genomic features. The numbers indicate the exact proportion (%). **d** NOR1-binding peaks at the promoters of *NADP-ME3*, *ARF5* and *AGL61*. **e** Expression levels of total *NOR1* (*NOR1*+*NOR2*+*NOR3*), *NADP-ME3*, *ARF5* and *AGL61* during fruit development in CG. **f** NOR1, NOR2 and NOR3 can activate the promoters of *NADP-ME3*, *ARF5* and *AGL61*. *P* values were determined by two-sided *t*-test. Data are given as mean ± s.d. with six biological replicates. **g** Expression levels of different *NOR1* copies in CG (*NOR1-allele-123*/*NOR1-allele-123*) and DSS (*NOR1-allele-1*/*NOR1-allele-1*) fruit. **h** Expression levels of different *NOR1* copies in YH (*NOR1-allele-123*/*NOR1-allele-123m*) mature fruit. Different letters indicate statistically significant differences (*P* < 0.05, one-way ANOVA with LSD test). **i** Proposed model of the expression activity of *NOR1* copies in different haplotypes. Data in **e**, **g** and **h** are given as mean ± s.d. with three biological replicates.

Interestingly, *NOR1* expression in cultivars carrying *NOR1-allele-123m* was lower than in cultivars with a comparable normalized *NOR1* depth (Fig. 4l). This may reflect diversification of the expression ability of different *NOR1* copies. To characterize the expression pattern of different *NOR1* copies, we used a specific sequence of each copy to quantify expression (Supplementary Fig. 20). In the *NOR1-allele-123* haplotype, *NOR1* expression was lower than that of *NOR2* and *NOR3*, whereas in *NOR1-allele-1*, *NOR1* was expressed at a high level (Fig. 5g), indicating that the gene expression level was diversified after duplication. Because H3K4me3 has a close correlation with gene expression^33^, we analyzed it at the *NOR1*, *NOR2*, and *NOR3* loci of *NOR1-allele-123* haplotype using the CUT&Tag method. Apparent modification signals at the *NOR1* locus were comparable to signals at *NOR2* and *NOR3* (Supplementary Fig. 21), indicating that the low expression level of *NOR1* in *NOR1-allele-123* is not due to the absence of H3K4me3 modification. In *NOR1-allele-123m*, *NOR3m* expression was almost undetectable (Fig. 5h), indicating that the large deletion disrupted the normal expression of *NOR3*. We lacked fruit from cultivars with the *NOR1-allele-12* haplotype and hence were unable to quantify the expression levels of different *NOR1* copies in *NOR1-allele-12*. Nevertheless, these results indicate that the expression levels of different *NOR1* copies were diversified, exemplifying for the diversified destiny of genes originating from recent duplication events (Fig. 5i).

### Selection of high-copy *NOR1* haplotypes has driven the development of modern early-maturing cultivars

Given that *NOR1-allele-123m* does not exhibit an apparent change in dosage compared with *NOR1-allele-1* (Fig. 5i), we eliminated accessions containing *NOR1-allele-123m* from analyses (Fig. 6a-f). Analysis of the normalized *NOR1* depth in different pear species^34^ revealed that the high-copy *NOR1* haplotypes were present in both wild and cultivated pears (Fig. 6a), and cultivated pears had a higher *NOR1* copy number than wild pears (Fig. 6b), suggesting the contribution of human selection. Further analysis of sand pear cultivars showed that improved cultivars had a higher *NOR1* copy number than landraces (Fig. 6c), indicating selection for the early-maturing trait in modern breeding programs. Sand pears can be classified into Chinese sand pears and Japanese sand pears. The present analysis indicated that, compared with Chinese sand pears, Japanese sand pears had a higher *NOR1* copy number (Fig. 6d). Further division of sand pears into landraces and improved cultivars revealed that the *NOR1* copy number was significantly higher in Japanese landraces than in Chinese landraces (Fig. 6e). However, no significant difference in *NOR1* copy number between Japanese and Chinese improved cultivars was detected (Fig. 6f), which may reflect comparable selection for the early-maturing trait in Chinese and Japanese breeding programs. Therefore, the ready availability of high-copy *NOR1* haplotypes among landraces and the market demand for early-maturing cultivars^35^ may have driven the breeding of early-maturing sand pears in Japan. To improve the quality of Chinese sand pears, Chinese breeders crossed Japanese sand pears with landraces to produce locally-adapted new cultivars while also selecting for the early-maturing trait. A kinship analysis of Chinese and Japanese sand pears partly reflected this breeding history (Fig. 6g). Analysis of normalized *NOR1* depth revealed that ‘Shinseiki’, owing to its higher *NOR1* copy number, was a crucial parent for improved Chinese sand pears with early to intermediate maturation (Fig. 6g), highlighting the important contribution of high-copy *NOR1* haplotypes to the development of modern pear cultivars. Given that China is the center of origin for pears, we propose that the *NOR1* duplication event may have first occurred in China and subsequently spread to Japan and other regions of the world. The introduction and hybridization of Japanese sand pears to improve Chinese cultivars since the twentieth century have introduced high-copy *NOR1* haplotypes into modern Chinese cultivars.

**Fig. 6.**
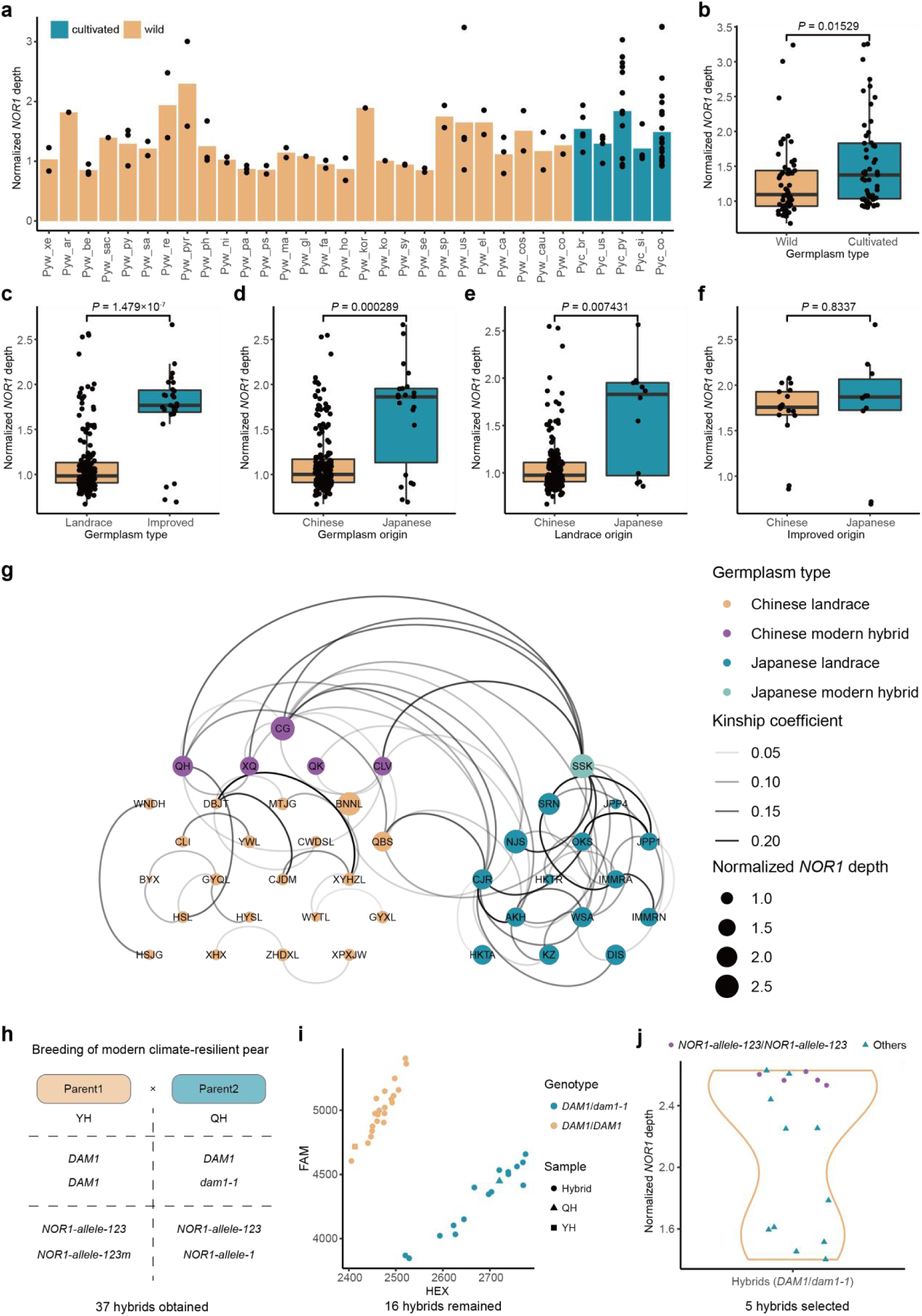
Selection of high-copy *NOR1* haplotypes has driven the development of modern early-maturing cultivars. **a** Normalized *NOR1* depth of 110 pear accessions from different species. The bars represent mean values of each group and each point represents one accession. The part of the group name before the underscore represents indicates wild or cultivated status, and the part after the underscore indicates the species. **b** Cultivated pears (*n* = 53) have higher normalized *NOR1* depth than wild pears (*n* = 57). **c** Improved sand pear cultivars (*n* = 27) have higher normalized *NOR1* depth than landraces of sand pear (*n* = 174). **d** Japanese sand pears (*n* = 22) have higher normalized *NOR1* depth than Chinese sand pears (*n* = 179). **e** Landraces of Japanese sand pear (*n* = 12) have higher normalized *NOR1* depth than landraces of Chinese sand pear (*n* = 162). **f** No significant difference was detected in normalized *NOR1* depth between improved sand pear cultivars from China (*n* = 17) and Japan (*n* = 10). **g** Kinship relationships estimated for Chinese and Japanese sand pears. **h** A hybrid population expected to breed early-maturing pear cultivars with low CR and early BD. **i** F_1_ hybrids carrying *dam1-1* were identified through KASP genotyping. **j** A violin plot showing the normalized *NOR1* depth of the F_1_ hybrids carrying *dam1-1*. Five accessions with the *NOR1-allele-1-2-3*/*NOR1-allele-1-2-3* genotype were selected. Accessions carrying *NOR1-allele-123m* were excluded from the analyses in **a**, **b**, **c**, **d**, **e** and **f**. *P* values in **b**, **c**, **d**, **e** and **f** were determined by two-sided *t*-test.

Based on our research findings, we have designed a breeding process that integrates genotypes for early BD, low CR, and short FDP (Fig. 6h) using YH (*DAM1*/*DAM1*, *NOR1-allele-123*/*NOR1-allele-123m*) and QH (*dam1-1*/*DAM1*, *NOR1-allele-1*/*NOR1-allele-123*) as the cross parents, both of which are modern cultivars with high quality fruit. This cross can yield extremely early-maturing individuals with climate resilience. In the newly generated F_1_ hybrid population (Fig. 6h), we filtered out 86% of non-target hybrids using a KASP marker for the *DAM1* locus and the read depth for the *NOR1* locus (Fig. 6i, j), thereby significantly improving the breeding efficiency.

## Discussion

The transition from a single reference genome to a pangenome represents a paradigm shift in our understanding of plant evolution^36^. The present study demonstrates that SVs, together with SNPs, collaboratively drive of phenotypic plasticity in *Pyrus*. While recent pangenomes in crops such as wheat^37^ and tomato^38^ have highlighted the role of SVs in grain hardness and fruit yield, the present findings on pear reveal a distinct landscape where SVs regulate developmental timing. Specifically, the pear genome has utilized two contrasting evolutionary mechanisms (loss-of-function via nonsense mutations and function enhancement via gene dosage expansion) to adapt to natural selection in a warm-winter environment and human selection for early fruit maturation.

Dormancy release is a critical survival strategy for temperate perennials. In peach (*Prunus persica*), the evergrowing phenotype arises from a large genomic deletion encompassing multiple *DAM* genes^39^. The present results indicate that low-chilling adaptation in sand pear landraces has evolved through convergent stop-gained variations in *DAM1*, which is more subtle. The independent origins of the *dam1-1* and *dam1-2* alleles suggest a strong selective sweep in southern subtropical regions of China, where maintenance of weak dormancy in a warm winter is essential for successful reproduction. Notably, two recent studies have focused on the contribution of genomic variations to climatic adaptation^40,41^, but these studies did not validate the impact of the variations on gene expression or function. The current study revealed that premature stop codons in *DAM1* may trigger nonsense-mediated decay^24^, resulting in the observed depletion of *DAM1* transcripts. This mechanism mirrors the SNP-triggered diversification in allele expression of the Arabidopsis *FLC* gene, which enhances the plasticity in plant environmental adaptability^42^, suggesting that modulating the dosage of MADS-box proteins is a universal strategy for phenological finetuning in annuals and perennials.

The regulation of fruit ripening has long been an important research focus in horticultural plants^43^. Our discovery of the *NOR1* CNV in pear introduces a novel dimension to the regulatory model of fruit ripening: SV-triggered change in gene dosage. Unlike the loss-of-function mutations in *DAM1*, the *NOR1* locus has undergone stepwise duplication (from one to three copies) during recent evolution, while the high copy-number haplotype has been selected and widely used in the breeding of early-maturing cultivars. We observed a negative correlation between *NOR1* dosage and the FDP, and all NOR1 copies can activate the expression of ripening-related genes, such as *NADP-ME3* and *ARF5*. These results suggest that the *NOR1* dosage is a critical factor accelerating fruit maturation. In the history of pear cultivation, breeders have selected for high-dosage alleles generated by CNVs to accelerate fruit ripening.

Stepwise duplication of the *NOR1* locus, which was dated to approximately 1.05–2.62 MYA, significantly predates human agriculture. This implies that the high-copy *NOR1* haplotypes were present as standing genetic variations in wild progenitor populations long before they were targeted by human selection. In a natural context, the high-dosage allele may have been disadvantageous in stable climates by decoupling flesh softening from optimal seed maturation. This restricts the allele frequency of such high-dose haplotypes. However, breeders impose strong directional selection on the high-copy *NOR1* haplotypes to drastically shorten the FDP. This scenario highlights a sophisticated model of human selection-mediated disruption of the allele frequency balance through fixing pre-existing SVs that were previously constrained by natural selection. This approach is similar to the selection of CNVs of *VRN-A1* in wheat^37^ and *MKK3* in barley^44^ to meet human needs.

The identification of causal SVs allows switching from phenotypic selection to haplotype-based design in breeding programs. Climate-resilient pear cultivars, which are capable of thriving in warm winters and producing high-quality fruit early in the season, can now be engineered by stacking the *dam1* (low CR and early BD) and *NOR1-allele-123* (short FDP) haplotypes. By utilizing the molecular markers developed in the present study, breeders can purge nontarget alleles at the seedling stage, reducing the breeding costs for this long-lived perennial. In conclusion, the pear pangenome resolves the genetic architecture of critical horticultural traits and provides a blueprint for adaptation of temperate fruit crops to a rapidly changing global climate.

## Methods

### Genome assembly

For the haploid consensus assemblies, Canu^45^ was used to correct the raw ONT reads and the corrected reads were assembled with SMARTdenovo^46^. Racon^47^ and Pilon^48^ were used to polish the raw contigs using long reads and short reads, respectively. Purge_haplotigs^49^ was used to remove the redundant sequences. LACHESIS^50^ was used to anchor the contigs at the chromosome level based on Hi-C reads. To assembled haplotype-resolved assemblies, hifiasm^51^ was used to assemble two haplotypes based on HiFi reads and Hi-C reads. The raw contigs were then anchored at the chromosome level using YaHS^52^.

### Genome annotation

For annotation of repeat sequences, a specific transposable element library for each assembly was constructed with the EDTA pipeline^53^. RepeatMasker^54^ was then used to annotate repeat sequences in each assembly. The insertion times of intact LTR-RTs were provided by EDTA using a mutation rate of 3.9 × 10^-9^ per site per year for apple^28^.

For annotation of gene models in the haploid consensus genomes, homology-based annotation, RNA-Seq-based annotation and de novo prediction were used to predict gene models. RNA-Seq-based annotation was obtained from TransDecoder (https://github.com/TransDecoder/TransDecoder), GeneMarkS-T^55^ and PASA^56^. GENSCAN^57^, GlimmerHMM^58^, Augustus^59^ and GeneID^60^ were used for de novo prediction. Homology-based annotation was generated by GeMoMa^61^. The annotations from different sources were merged using EVM^62^. For the haplotype-resolved genomes, we first used the BRAKER3 pipeline^63^, which utilized proteins and RNA-Seq data to obtain the Augustus annotation. We then used the MAKER^64^ pipeline to integrate the evidence to obtain high-quality gene models.

### Pan-genome analyses

We clustered proteins from the 11 pear genomes assembled in this study and 13 publicly available pear genomes^15,65,66,67,68,69,70,71^ using OrthoFinder^72^. We designated the gene clusters as core (present in all 24 genomes), shell (present in 2–23 genomes) and private (present in one genome). The genes in the core and shell clusters were core and shell genes, respectively. The genes in the private cluster and those that failed to be clustered were designated as private genes. The *Ka*/*Ks* values were calculated with TBtools^73^. For the assembly-based SV detection, query assemblies were aligned to the YL genome using minimap2^74^. SVIM-asm^75^ was then used to detect SVs. The PAVs were merged using Jasmine^76^. The vg toolkit^77^ was used to construct a graph-based pangenome based on PAVs. For the genotyping of PAVs in sand pear, we merged PAVs from 11 sand pear-related assemblies using Jasmine^76^ and genotyped them using Paragraph^78^ with a graph-based method.

### Population sequencing data analyses

Fastp^79^ was used to generate clean short reads. Clean short reads were aligned to the reference genome (YL) using BWA^80^. Picard (https://broadinstitute.github.io/picard/) was used to mark duplicated reads. BCFtools^81^ was used to call SNPs and InDels. Bi-allelic SNPs with minor allele frequency > 5% and missing rate < 20% were used for further analyses. Nucleotide diversity (π) and genetic distance (*F*_st_) were calculated with VCFtools (https://vcftools.github.io/index.html). PLINK^82^ was used to perform the principal component analysis and kinship calculation. FastTree^83^ was used to construct the phylogenetic tree which was visualized using iTOL^84^. Independent SNPs were input to ADMIXTURE^85^ to do population structure analyses. GAPIT3^86^ was used to perform GWAS based on SNP and InDel markers. The kinship of different accessions was calculated by PLINK2^82^.

### Gene expression quantification

In addition to the data generated in this study, public RNA-Seq data^87,88,89,90,91,92,93^ were also used for analyses. For the RNA-Seq data with short reads, we used HISAT2^94^ to perform the alignment and featureCounts to count reads. The expression level was presented as transcripts per million (TPM). For the expression quantification of two alleles in QH, we used Liftoff^95^ to identify the allelic gene pairs in two haploids. The two haploids were merged to generate a diploid genome. Minimap2^74^ was used to align the full-length RNA-Seq data to the diploid genome, retaining only the best alignments. FeatureCounts^96^ was used to count reads for each gene. We obtained normalized read counts by dividing the raw read counts by the total number of assigned reads. To analyze the association between the PAV genotype and the gene expression level, we identified PAVs within 5000 bp upstream and downstream of each gene, and used the linear regression model to analyze the association between the PAV genotype and the corresponding gene expression level.

### Chilling requirement evaluation

One-year-old shoots from different cultivars were collected from Changxing County of Zhejiang Province on December 19, 2025. The temperature at the site where the cultivars were grown was monitored hourly. The collected shoots were placed in a green-house under the forcing condition (22 °C/20 °C, 10-h day/14-h night cycle) for 28 days. When the bud breaking rate exceeded 50%, it indicated that the required chilling had been satisfied and dormancy had been released.

### DNA extraction and DNA-qPCR

Fresh leaves were frozen with liquid nitrogen and ground into powder. Approximately 0.1–0.2 g powder was added to 700 µL of cetyltrimethylammonium bromide solution preheated to 65°C and mixed with 20 µL β-mercaptoethanol. The lysate was incubated at 65°C for 30 min, then 700 μL chloroform/isoamyl alcohol (24:1, v/v) was added and mixed for 10 min. After centrifugation for 10 min at 12,000 rpm, 400 µL of the supernatant was transferred to a new centrifuge tube. Isopropanol (400 µL) was added to the centrifuge tube and allowed to stand at –20 °C for 30 min. The mixture was centrifuged at 12,000 rpm for 10 min and the supernatant was discarded. Next, 500 µL of 75% ethanol was added to the tube and the mixture was centrifuged at 13,000 rpm for 5 min. After discarding the supernatant, the precipitate was dissolved in ddH_2_O to yield genomic DNA. The DNA-qPCR analysis amplified the genomic DNA as the template, and the other steps were similar to the method previously described for quantitative real-time PCR^67^. A single-copy gene *EXPA7* was used for standardization. The primers used for DNA-qPCR are listed in Supplementary Table 5.

### Calculation of normalized *NOR1* depth

The clean reads were aligned to the YL genome which contained only one *NOR1* copy, using BWA^80^. Picard (https://broadinstitute.github.io/picard/) was used to mark duplicated reads in the BAM files. PanDepth^97^. was used to calculate the mean depth of the whole genome and the mean depth of the *NOR1* region. The mean depth of the *NOR1* region was divided by the mean depth of the whole genome to derive the normalized *NOR1* depth.

### Phylogenetic tree construction for NOR1 and related proteins

The protein sequence of pear NOR1 was used as the query to search for homologous proteins in *Arabidopsis* and tomato using BLASTP. The NOR protein from tomato was used as the query to search for homologous proteins in pear. The top five hits from the BLASTP results were selected. The proteins were aligned using MUSCLE^98^. A phylogenetic tree was constructed with IQTREE^99^ based on the maximum-likelihood method and visualized using iTOL^84^.

### DAP-Seq analysis

DAP-Seq was performed according to a previously described method^100^. The correct DAP-seq library and input libraries were sequenced on an Illumina NovaSeq 6000 platform to produce PE150 reads. The clean reads were aligned to the reference genome using BWA^80^. Picard (https://broadinstitute.github.io/picard/) was used to mark duplicated reads. MACS2^101^ was used to detect peaks. Peaks from two replicates were merged using IDR^102^. We used MEME-ChIP (https://meme-suite.org/meme/tools/meme-chip) to identify conserved motifs. ChIPseeker^103^ was used to annotate the distance from a peak to the transcription start site. The BAM files were converted to bigWig files for visualization using deepTools^104^.

### Dual-luciferase assay

We performed dual-luciferase assays following our previously described method^105^. The full-length CDS of each candidate gene was cloned into the BamHI/HindIII site of the pGreenII 0029 62-SK vector and the candidate promoters were cloned into the NcoI/HindIII site of the pGreenII 0800-LUC vector. All vectors were transformed into *Agrobacterium tumefaciens* strain GV3101 cells harboring the pSoup vector. These two *A*. *tumefaciens* strain GV3101 cultures were merged at a ratio of SK:LUC = 1:10 and then injected into *Nicotiana benthamiana* leaves. After incubation for 48 hours, the firefly luciferase and Renilla luciferase activities were measured using the Dual-luciferase Reporter Assay System (Promega, USA). The primers used for constructing the vectors are listed in Supplementary Table 5.

### Validation of stop-gained SNPs in the *DAM1* locus and KASP genotyping

The DNA template was diluted to approximately 100 ng/µL. PCR products were generated by amplifying the sequences surrounding the SNPs and Sanger sequencing was performed on these PCR products to determine the presence of the corresponding SNPs based on the sequencing signals. For KASP genotyping, specific amplification primers were first designed, and the template was amplified using the PCR mix for KASP genotyping. After the reaction was completed, the fluorescence intensities at two different wavelengths in the reaction mixture were measured to determine the genotype of the corresponding base.The primers used for genotyping are listed in Supplementary Table 5.

## Acknowledgements

We thank Prof. Yuanwen Teng (Zhejiang University) for valuable discussions and suggestions and Mr. Kunfeng Li (Zhejiang University) for his assistance in tree cultivation. This research was supported by the National Natural Science Foundation of China (32272639 and 32472702 to S.B., 32302466 to Jia W.), Specialized Fund for New Agricultural Cultivar Breeding in Zhejiang Province (2021C02066-5 to D.C.), Science and Technology Promotion for Agriculture Project of Shanghai (2019-1-3 to S.J.), Titled Research and Demonstration of Green and Efficient Production Technology in Pear Orchards (sponsored by Shanghai Municipal Agricultural Commission to S.J.), the Earmarked Fund for the China Agriculture Research System (CARS-28-03 to D.C.), and China Postdoctoral Science Foundation (2024M762852 to Y.G.).

## Author contributions

S.B., D.Z., X.Z., D.C. and S.J. conceived and designed the project. Y.G. performed bioinfomatics analyses and other major works described in the article. W.W., Y.L. and L.W. performed experiments. Jiahao W., M.D., C.W., L.T., H.L. and S.J. collected and provided plant materials. Y.G. and S.B. drafted the manuscript. Jia W., C.J., J.S., H.X. and J.N. revised the manuscript.

## Competing interests

The authors declare no competing interests.

## Notes

### Competing Interest Statement

The authors have declared no competing interest.

