## Supplementary Note for "A graph-based pangenome reveals the genetic basis of climate-resilient and horticultural traits in pear"

**Supplementary Note 1. Introgression and diversification between cultivated and wild pear species**

To analyze the introgression between cultivated and wild pears, we collected sequencing data for 70 wild pear accessions (including 30 *P. pashia*, 25 *P. xerophila* and 15 *P. ussuriensis*) and 33 cultivated pear accessions (all of them are *P. pyrifolia*). These wild pears were native to different regions of China: *P. pashia* from southwestern China, *P. xerophila* from northwestern China and *P. ussurensis* from northeastern China. The *P. pyrifolia* cultivars originated from China and Japan. The SNP-based phylogenetic tree showed that the individuals of *P. pyrifolia*, *P. pashia*, *P. ussuriensis*, and *P. xerophila* clustered together within their respective species (Supplementary Fig. 1a). It is noted that *P. pyrifolia* accessions formed multiple branches (Supplementary Fig. 1a), which may be due to the wide range of geographic origins of these accessions. Population structure of these pear accessions was analyzed. When K equaled the optimal number (K = 4), the results successfully distinguished different species (Supplementary Fig. 1b). Notably, the population structure result indicated the introgression from *P. pyrifolia* to *P. ussuriensis* and *P. xerophila*, as well as from *P. pashia* to *P. pyrifolia* (Supplementary Fig. 1b). These introgressions may arise from geographic overlap among different species.

Combining the results of population structure, phylogenetic tree, *D*-statistic and principle component analysis (Supplementary Fig. 1c), we deduced that ML likely belongs to *P. pyrifolia* whereas YNSL has a genetic background derived from both *P. pyrifolia* and *P. pashia*. We detected that NH harbored a larger proportion of gene pools from *P. pyrifolia* (Supplementary Fig. 1b), which was also validated by the *D*-statistic (Supplementary Fig. 1d).

To analyze the genetic diversity and divergence of different *Pyrus* species, we calculated nucleotide diversity ($\pi$) and genetic distance ($F_{st}$). Interestingly, we found that *P. pyrifolia*, as a cultivated species, exhibited the highest genetic diversity, followed by *P. xerophila*, *P. pashia*, and *P. ussuriensis* (Supplementary Fig. 1e). This may be attributed to the wide distributions of *P. pyrifolia* accessions. The $F_{st}$ results indicated that the differentiation between *P. pyrifolia* and *P. xerophila* was the lowest (Supplementary Fig. 1f), suggesting that they have relatively close genetic backgrounds. *P. pashia* and *P. ussuriensis* exhibit the highest differentiation, which may be due to their geographically distant distributions and lack of gene flow.
