## Supplementary Fig. for "A graph-based pangenome reveals the genetic basis of climate-resilient and horticultural traits in pear"

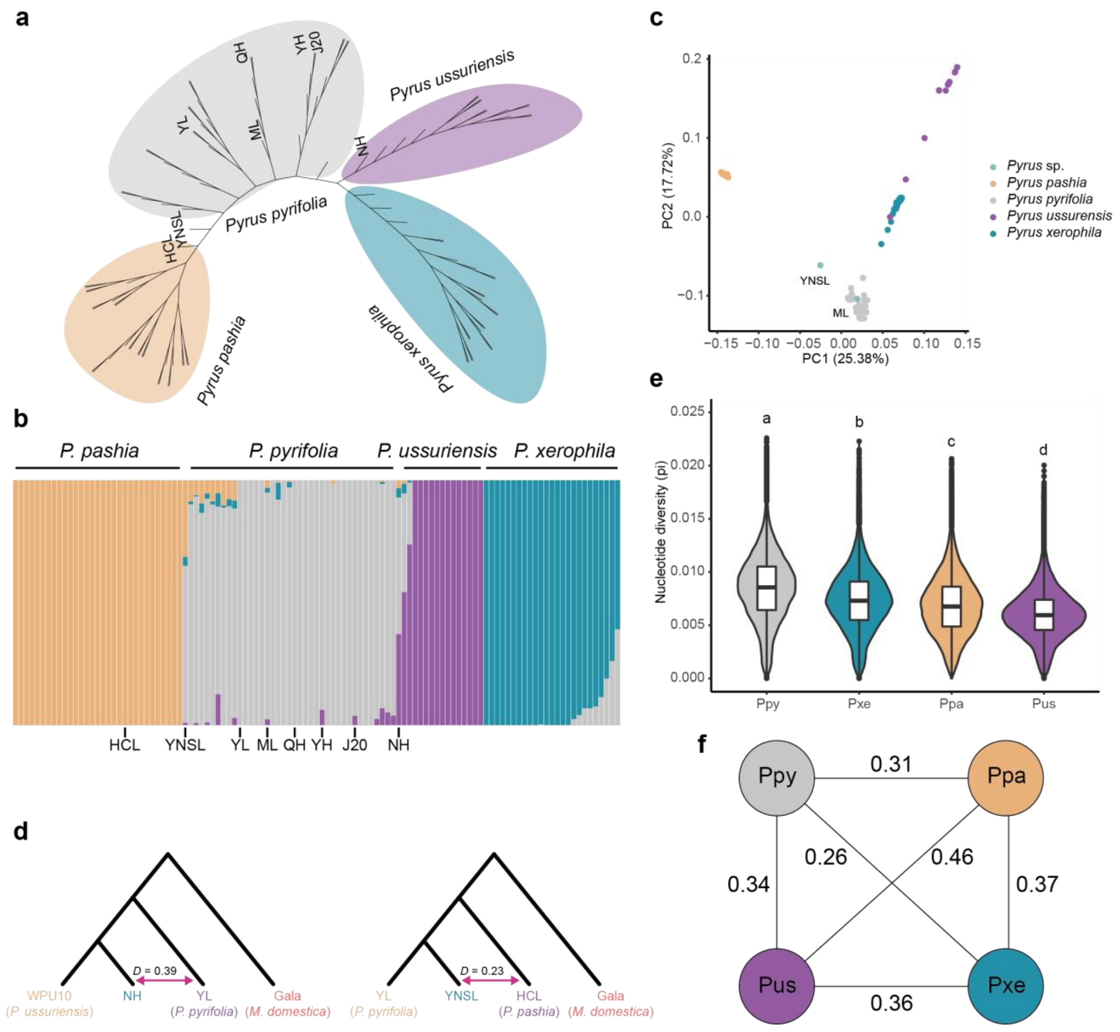


**Supplementary Figure 1. Introgression and diversification between different pear species. a** Phylogenetic tree of 103 pear accessions from different species. **b** Model based clustering of 103 pear accessions from different species when the number of ancestry kinship is 4. **b** Principle component analysis of 103 pear accessions based on SNPs. **d** *D*-statistic showed NH and YNSL may originated from interspecific hybridization. **e** Nucleotide diversity (π) of different pear species. Different letters indicate statistically significant differences ($P < 0.05$, one-way ANOVA with least significant difference test). **f** Population divergence ($F_{st}$) between different pear species. Ppy: *P. pyrifolia*, Pus: *P. ussuriensis*, Ppa: *P. pashia*, Pxe: *P. xerophila*.

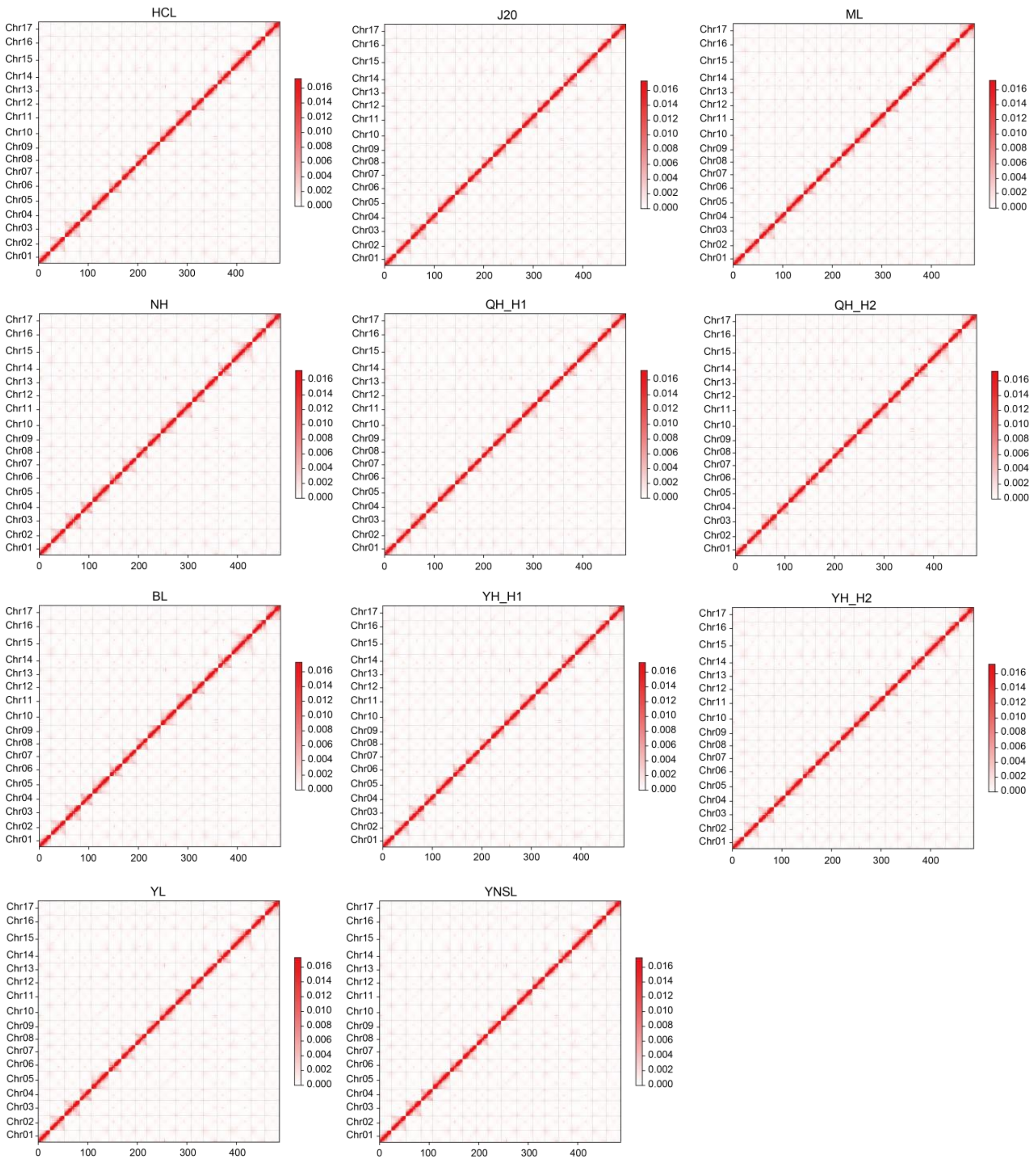


**Supplementary Figure 2. Hi-C contact map of 11 genomes.** The heatmaps indicate the KR normalized counts with a bin size of 500 Kb.

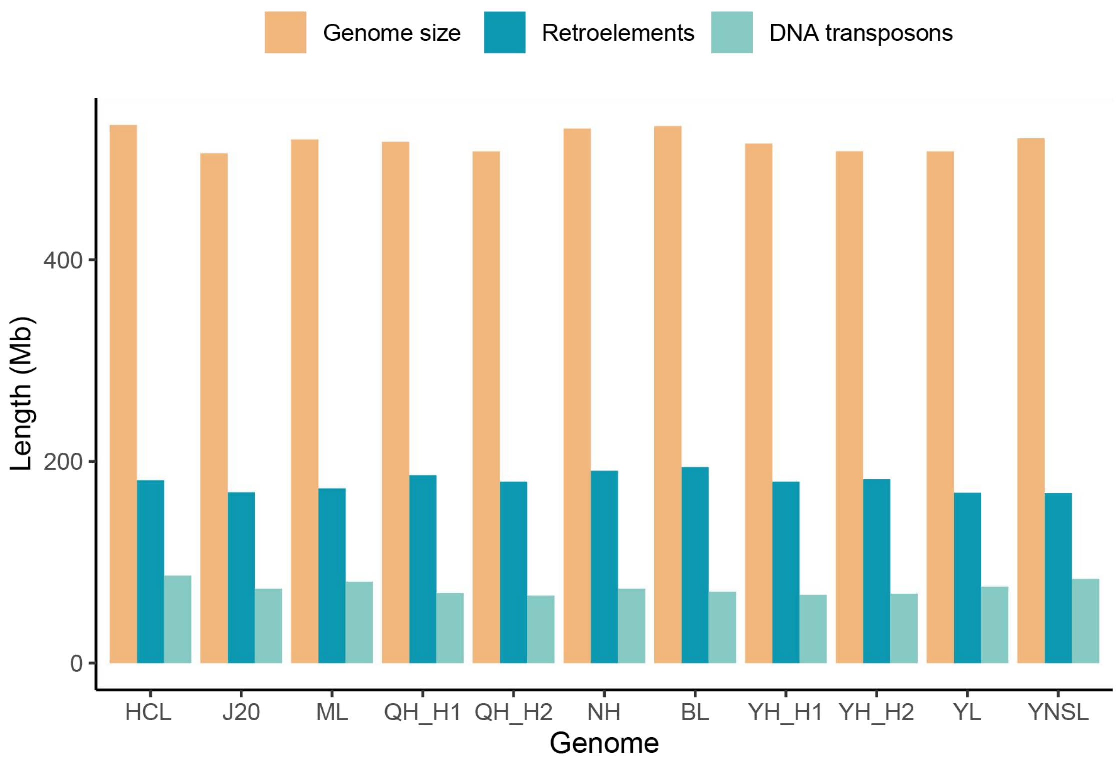


**Supplementary Figure 3. Base length of retroelements and DNA transposons in each genome.**

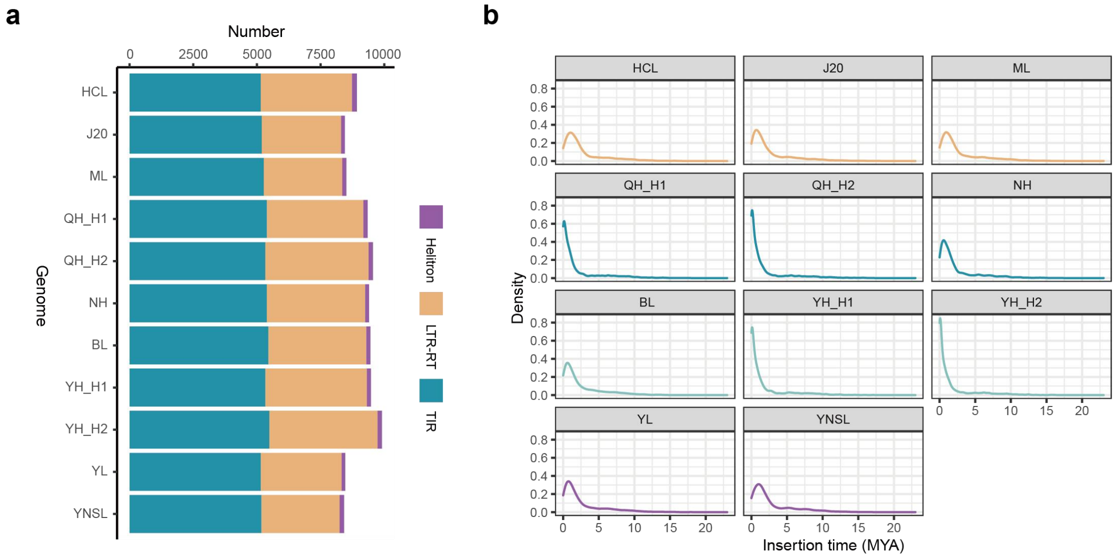


**Supplementary Figure 4. Intact transposable elements (TEs) in each genome. a** Number of intact TEs in each genome. **b** Insertion time of intact LTR-RTs in each genome.

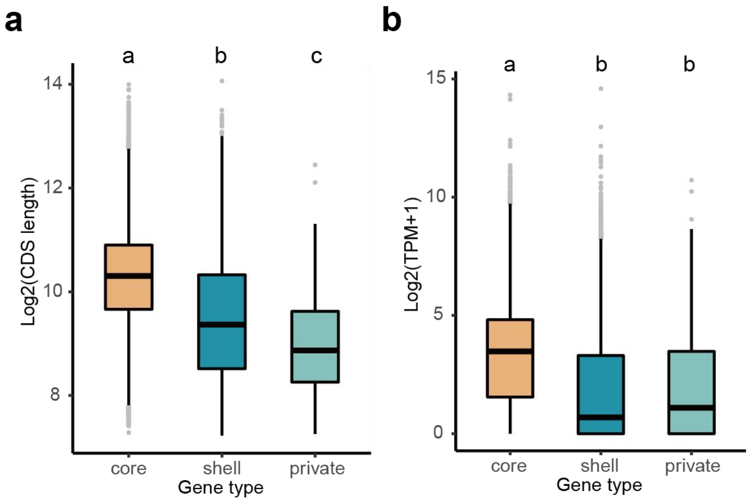


**Supplementary Figure 5. CDS length (a) and expression levels (b) of core, shell and private genes.** Different letters indicate statistically significant differences ($P < 0.05$, one-way ANOVA with LSD test). In boxplots, the 25% and 75% quartiles are shown as the lower and upper edges of the boxes, respectively, and central lines denote the median. The whiskers extend to 1.5× the interquartile range.

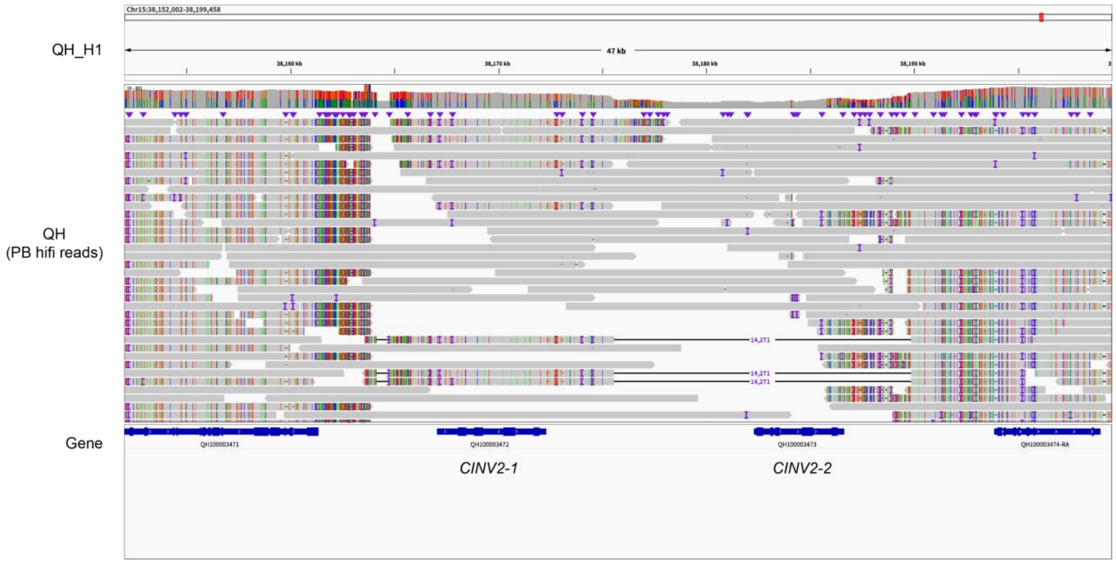


**Supplementary Figure 6. Read alignments showing a large deletion resulting in loss of *CINV2-2*.**

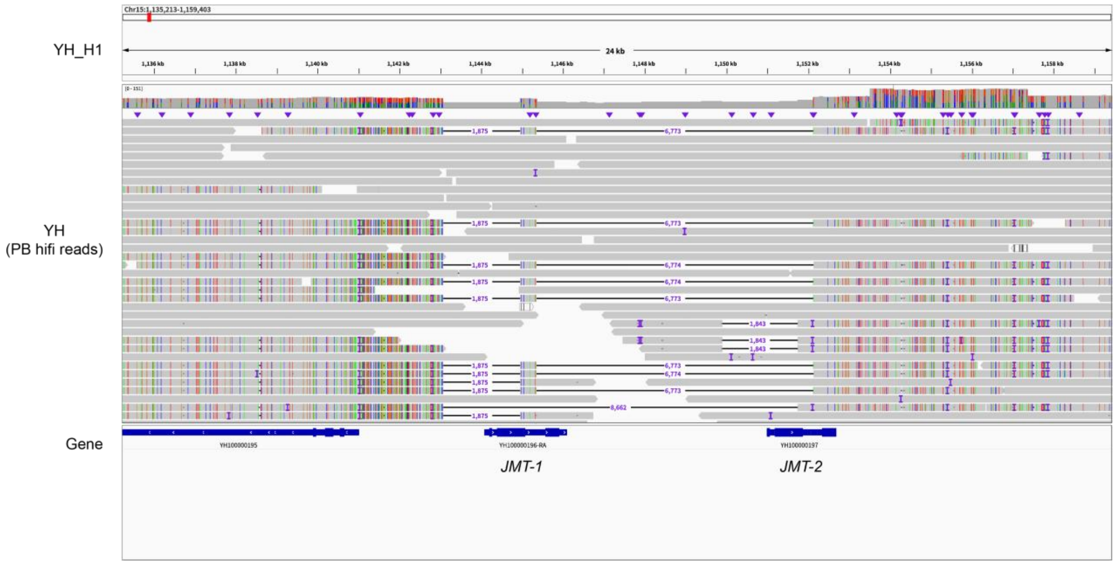


**Supplementary Figure 7. Read alignment showing large deletions resulting in loss of CDSs of *JMT-1* and *JMT-2*.**

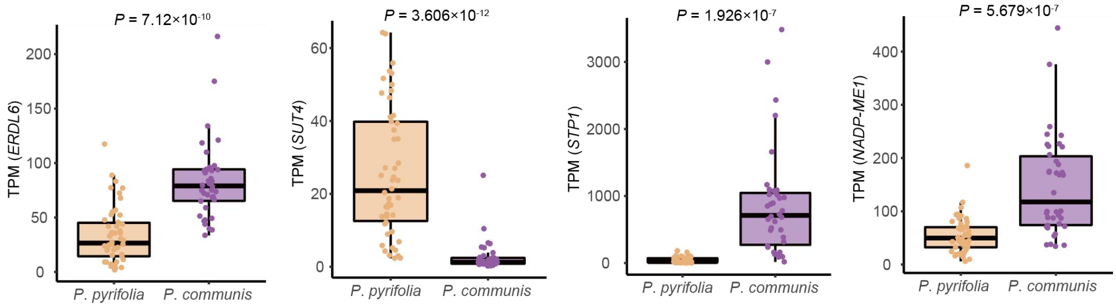


**Supplementary Figure 8. Sugar and acid related genes that differentially expressed between *P. pyrifolia* and *P. communis*.** *P* values were determined by two-sided *t*-test.

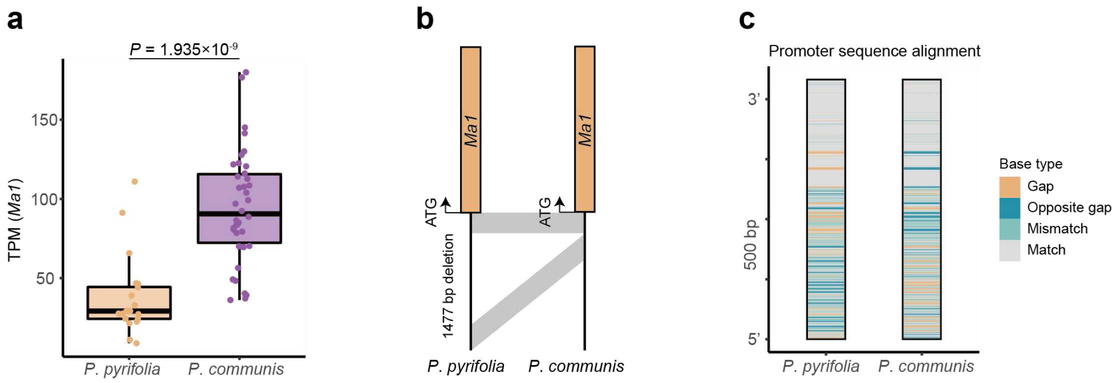


**Supplementary Figure 9. Difference between *Ma1* promoters of *P. pyrifolia* and *P. communis*.** **a** Differences between *P. pyrifolia* (accessions that do not carry the PAV causing *Ma1* loss) and *P. communis*. **b** A PAV in the *Ma1* promoter distinguishes *P. pyrifolia* from *P. communis*. **c** Sequence alignment of *Ma1* promoters from *P. pyrifolia* and *P. communis*.

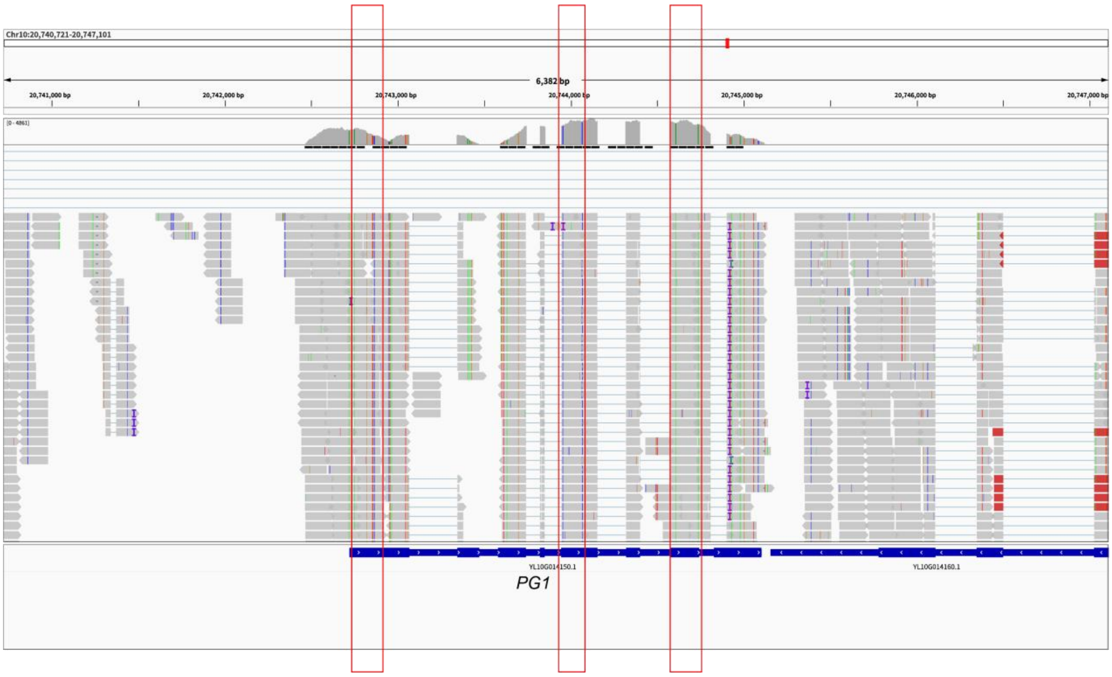


**Supplementary Figure 10. Allele-specific expression of *PG1* in a interspecific hybrid of *P. pyrifolia* and *P. communis* (fruit RNA).** Red rectangles highlight example SNPs specific to the *P. communis* parent.

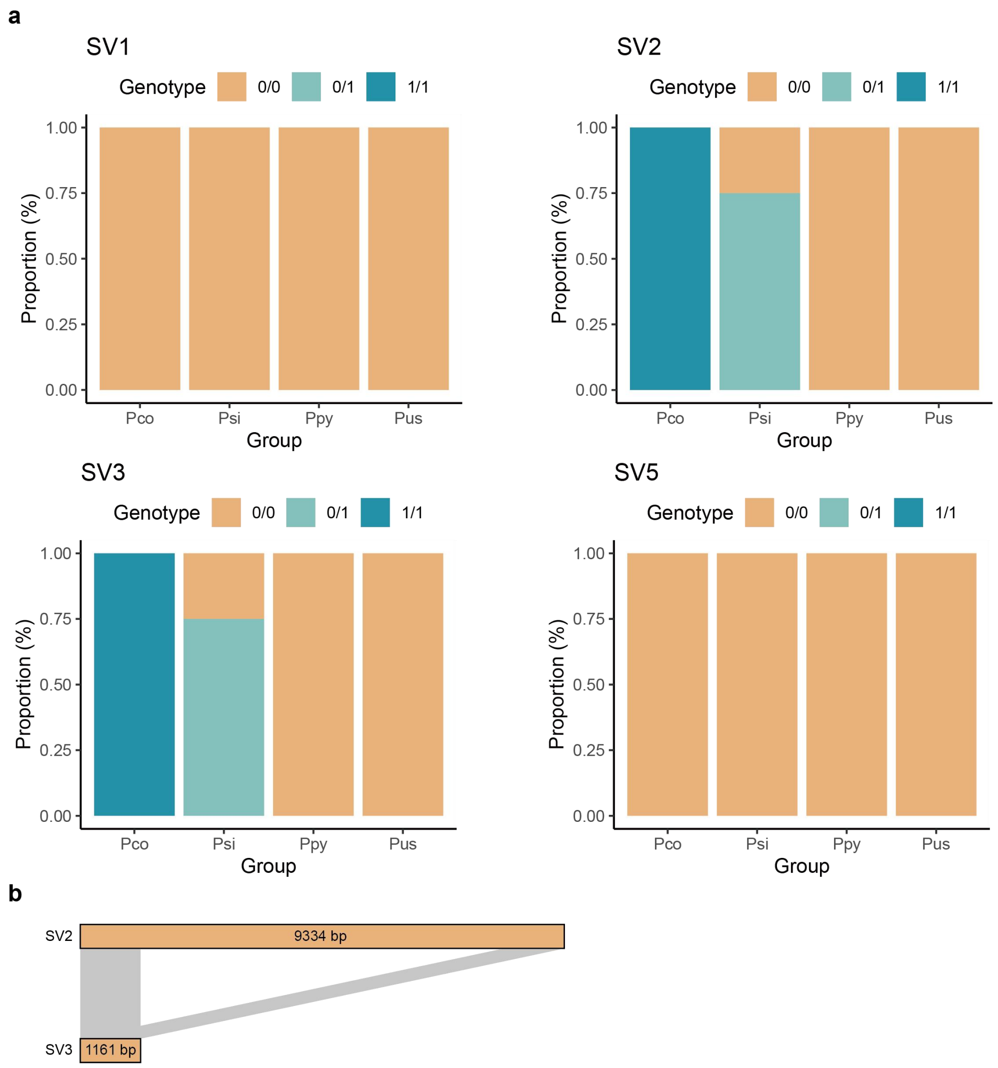


**Supplementary Figure 11. Genotyping results of SVs at *PG1* locus. a** genotyping results of SV1, SV2, SV3 and SV5. Pco: *Pyrus communis*, Psi: *Pyrus sinkiangensis*, Ppy: *Pyrus pyrifolia*, Pus: *Pyrus ussuriensis*. **b** The sequence of SV3 is similar to the sequences at both ends of SV2, which may explain why genotyping results for these two SVs are identical when using short reads.

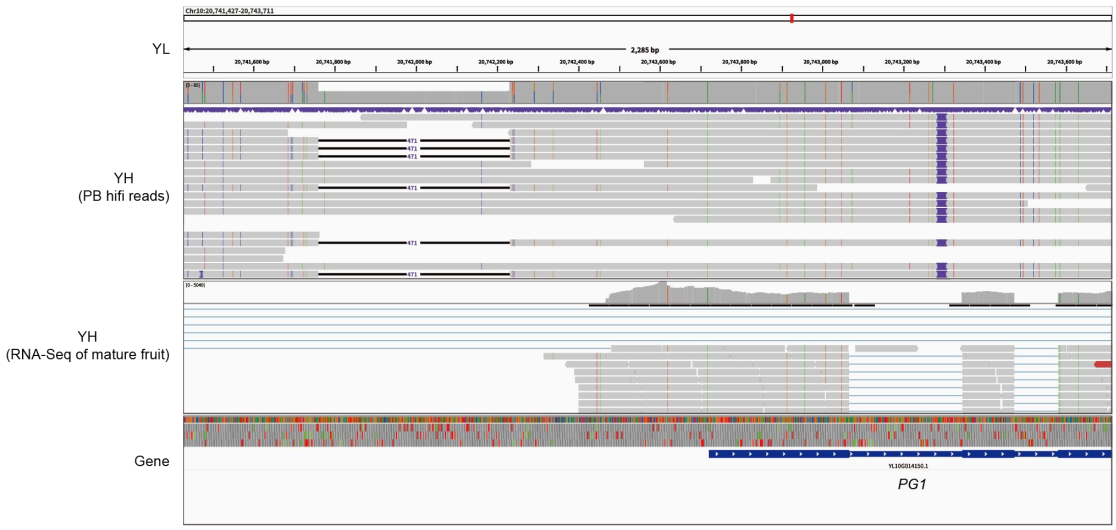


**Supplementary Figure 12. Allele-specific expression of *PG1* in mature fruit of YH.**

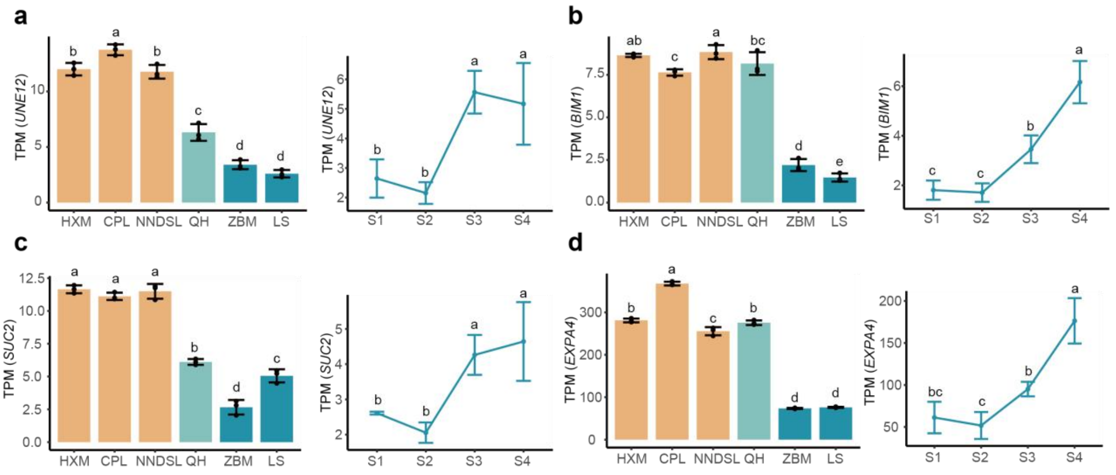


**Supplementary Figure 13. Expression levels of genes highly negatively correlated with *DAM1* in pear buds.** The horizontal axes represent different accessions or sampling times of same cultivar. Different letters indicate statistically significant differences ($P < 0.05$, one-way ANOVA with LSD test). Data are given as mean ± s.d. with three biological replicates.

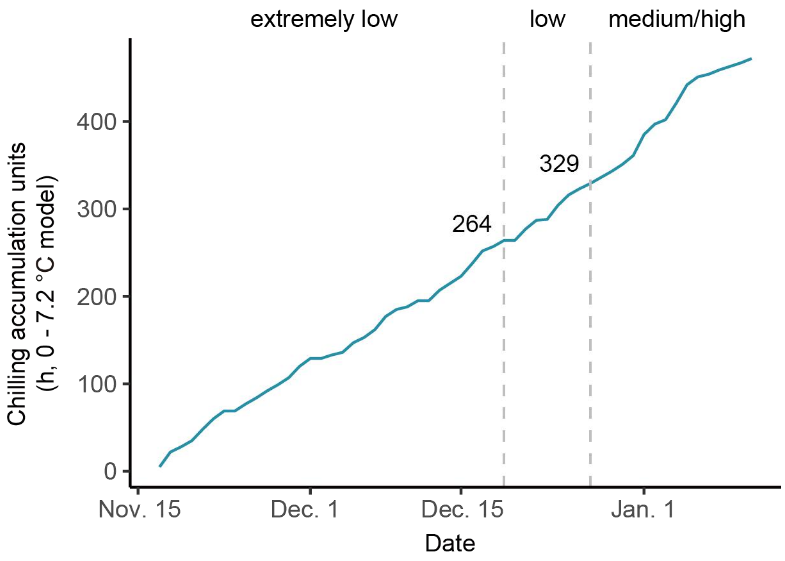


**Supplementary Figure 14. Chilling accumulation in Changxing County, Zhejiang Province, China.** The vertical lines indicate the dates of shoot sampling (Dec. 19, 2025 and Dec. 27, 2025).

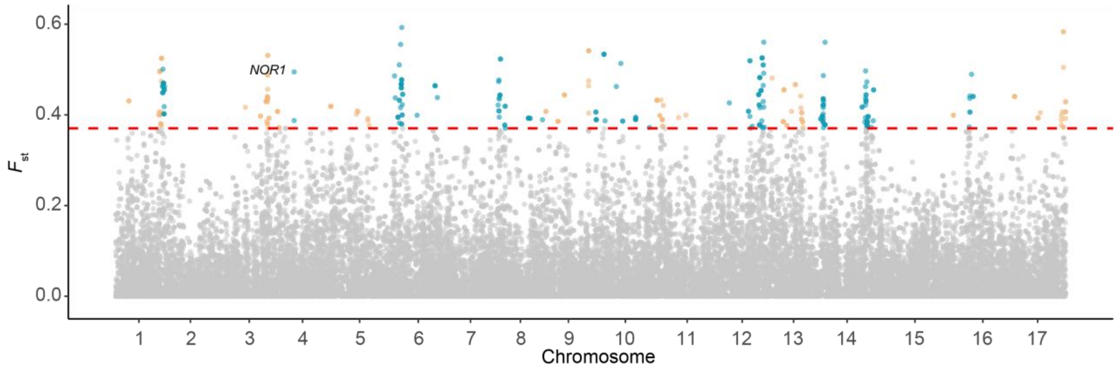


**Supplementary Figure 15. Selection signatures ($F_{st}$) in a pear population based on PAVs.** The red dashed line indicates the top 1% threshold.

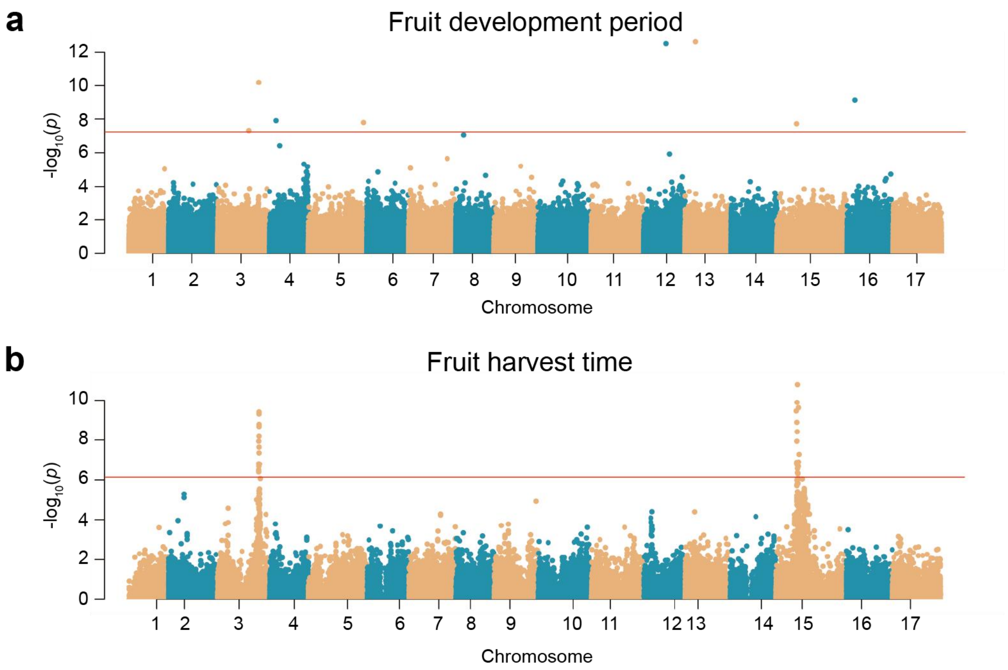


**Supplementary Figure 16. Genome-wide association study results for fruit development period (a) and fruit harvest time (b).** In **a**, InDels were used as markers with the FarmCPU model. In **b**, SNPs were used as markers with the mixed linear model. The red lines in **a** and **b** indicate the significant cutoff calculated as $0.05/n$, where $n$ is the total marker number.

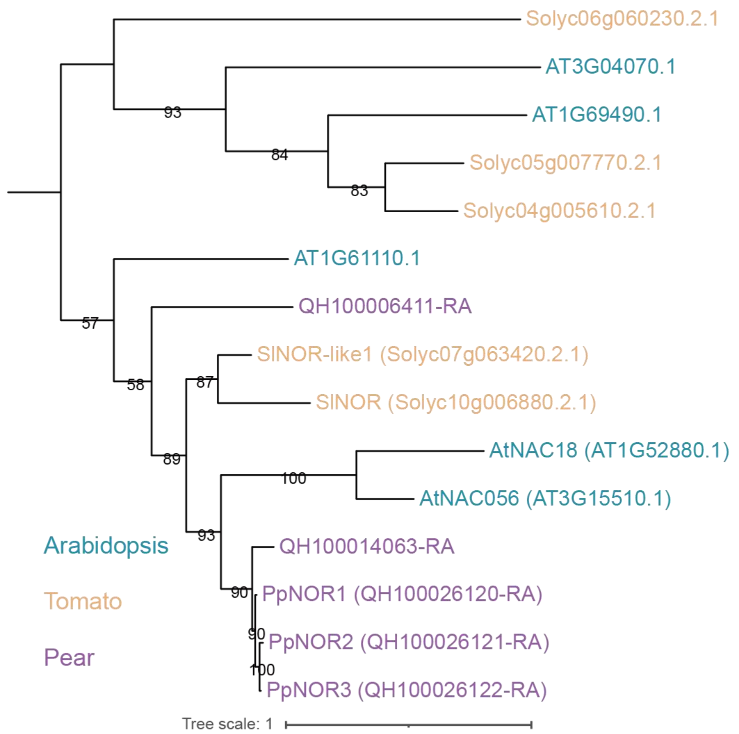


**Supplementary Figure 17. Phylogenetic tree of pear *NOR1* copies and their homologs in pear, *Arabidopsis* and tomato.** The tree was inferred using the maximum-likelihood method. Numbers at the branches represent ultrafast bootstrap values from 1,000 replicates

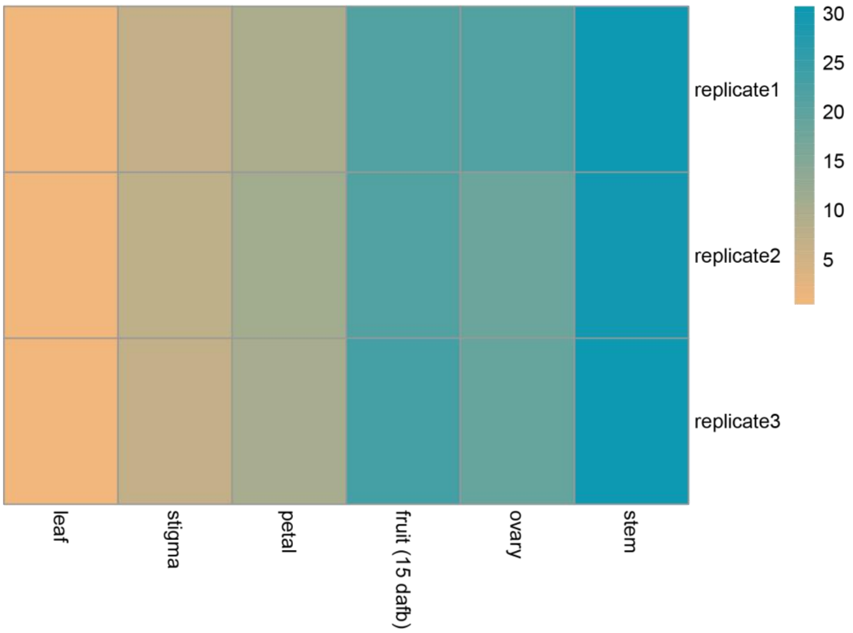


**Supplementary Figure 18. A heatmap showing *NOR1* expression levels in different pear tissues.** TPM values were used to represent expression levels.

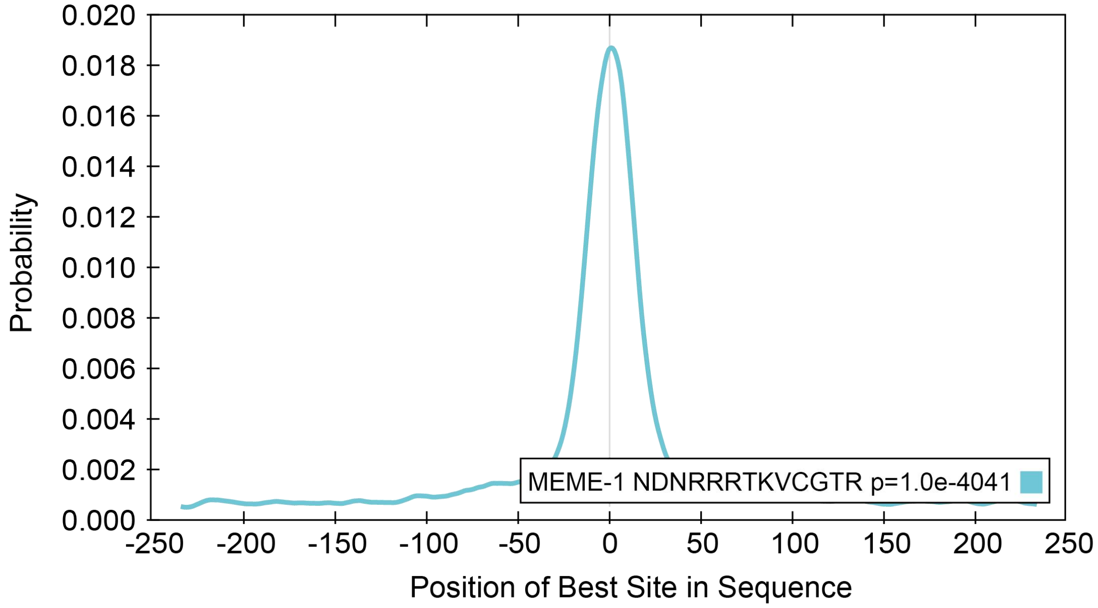




**Supplementary Figure 19. Local motif enrichment analysis for the motif discovered in DAP-Seq experiment.** The 500 bp sequences surrounding summits of peaks were used for the analysis.

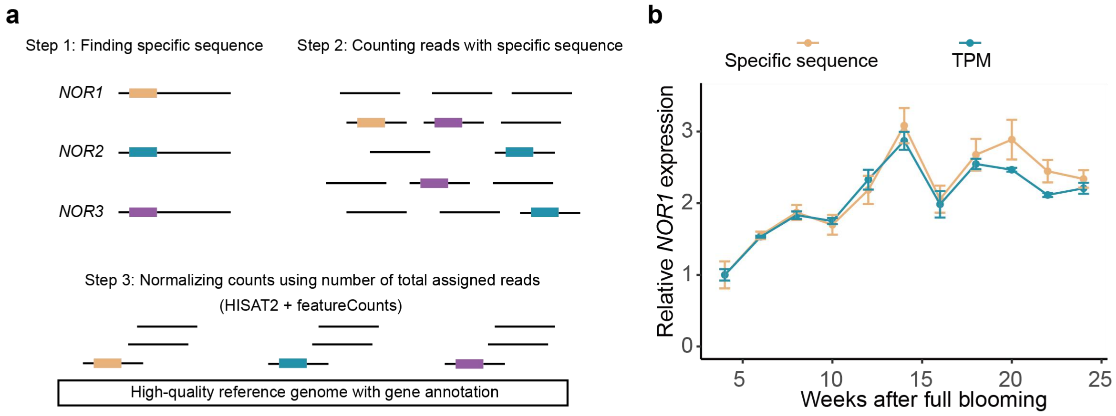


**Supplementary Figure 20. The pipeline used to calculate the expression levels of different *NOR1* copies. a** detailed steps of this pipeline. **b** Consistency between relative *NOR1* expression calculated by this pipeline (using specific sequence) and TPM in fruit of DSS (*NOR1-allele-1*/*NOR1-allele-1*). Data are given as mean ± s.d. with three biological replicates.

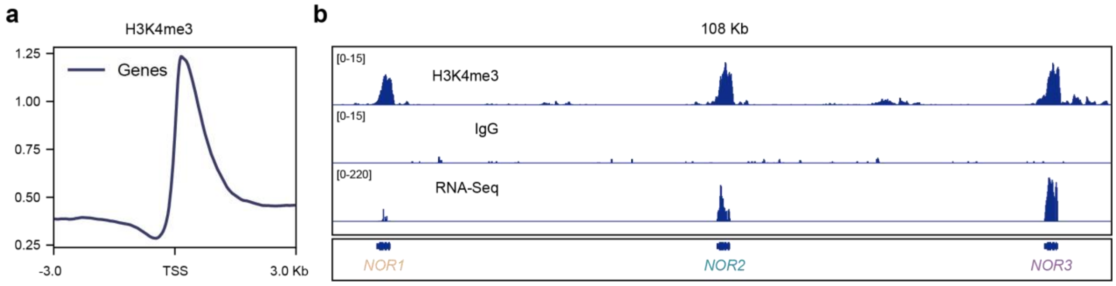


**Supplementary Figure 21. H3K4me3 modification at different *NOR1* copies.**

**a** Distribution of H3K4me3 modification signals relative to the transcription start site. **b** H3K4me3 modification and RNA-Seq signals at gene bodies of *NOR1*, *NOR2* and *NOR3*.
