## Supplementary Table for "A graph-based pangenome reveals the genetic basis of climate-resilient and horticultural traits in pear"

**Supplementary Table 1. Sequencing reads for genome assembling**

| Accession | Species | Platform of long reads | Total bases of long reads | Total bases of short reads | Hi-C bases |
|---|---|---|---|---|---|
| YL | *Pyrus pyrifolia* | ONT | 81,776,548,977 | 25,531,443,022 | 51,061,536,220 |
| ML | *Pyrus* sp. | ONT | 61,194,995,767 | 30,019,485,612 | 52,687,347,916 |
| J20 | *Pyrus pyrifolia* | ONT | 60,924,378,499 | 25,230,185,128 | 50,541,747,346 |
| NH | *Pyrus ussuriensis* | ONT | 61,617,815,956 | 28,851,737,326 | 52,562,488,006 |
| HCL | *Pyrus pashia* | ONT | 25,979,477,580 | 28,164,789,342 | 61,127,125,308 |
| BL | *Pyrus communis* | ONT | 62,807,754,405 | 29,493,643,954 | 50,331,541,192 |
| YNSL | *Pyrus* sp. | ONT | 60,534,025,393 | 29,175,224,656 | 55,337,551,048 |
| QH | *Pyrus pyrifolia* | PacBio HiFi | 29,672,748,306 | 25,052,352,900 | 35,449,986,300 |
| YH | *Pyrus pyrifolia* | PacBio HiFi | 38,000,122,813 | 28,527,250,800 | 35,775,705,300 |

**Supplementary Table 2. Quality assessment of 11 pear genomes**

| Genome | Type | Base | Base (without N) | Scaffold N50 | Contig N50 | LAI | BUSCO (genome) | Gene number | BUSCO (protein) |
|---|---|---|---|---|---|---|---|---|---|
| YL | haploid consensus | 507,439,638 | 507,412,238 | 27,913,133 | 2,230,012 | 20.91 | 97.4 | 40,891 | 93.8 |
| ML | haploid consensus | 519,267,277 | 519,234,377 | 28,263,737 | 1,961,850 | 19.19 | 98.0 | 41,837 | 94.9 |
| J20 | haploid consensus | 505,476,180 | 505,451,880 | 27,986,799 | 2,880,971 | 20.34 | 98.3 | 41,221 | 95.9 |
| NH | haploid consensus | 529,935,098 | 529,912,298 | 30,085,510 | 2,828,000 | 21.17 | 98.3 | 42,350 | 95.4 |
| HCL | haploid consensus | 533,554,455 | 533,523,855 | 29,806,954 | 2,386,832 | 19.89 | 98.6 | 41,135 | 96.2 |
| BL | haploid consensus | 532,506,626 | 532,474,626 | 29,789,823 | 2,033,013 | 21.52 | 98.2 | 42,053 | 95.5 |
| YNSL | haploid consensus | 520,343,174 | 520,312,974 | 29,948,109 | 2,346,863 | 18.31 | 97.9 | 42,613 | 95.2 |
| QH_H1 | phased | 516,854,776 | 516,854,176 | 28,858,000 | 26,492,385 | 20.22 | 99.0 | 42,846 | 98.0 |
| QH_H2 | phased | 507,420,565 | 507,419,365 | 29,349,252 | 23,406,094 | 23.54 | 99.0 | 46,357 | 97.9 |
| YH_H1 | phased | 515,175,365 | 515,174,865 | 28,957,274 | 26,362,166 | 22.75 | 98.8 | 45,214 | 97.6 |
| YH_H2 | phased | 507,527,487 | 507,526,987 | 28,889,821 | 28,836,007 | 23.91 | 99.0 | 44,477 | 97.9 |

**Supplementary Table 3. Repeat annotation of 11 pear genomes**

| Genome | Total bases | Repeat (%) | Retroelement (%) | DNA transposon (%) | Unclassified (%) | Simple repeat (%) | Low complexity (%) |
|---|---|---|---|---|---|---|---|
| YL | 507,439,638 | 51.27 | 33.27 | 14.94 | 1.94 | 0.90 | 0.21 |
| ML | 519,267,277 | 52.00 | 33.37 | 15.53 | 1.98 | 0.90 | 0.22 |
| J20 | 505,476,180 | 51.23 | 33.50 | 14.58 | 1.99 | 0.93 | 0.23 |
| NH | 529,935,098 | 52.84 | 35.97 | 13.93 | 1.73 | 0.99 | 0.22 |
| HCL | 533,554,455 | 53.31 | 34.00 | 16.24 | 1.89 | 0.97 | 0.22 |
| BL | 532,506,626 | 52.68 | 36.47 | 13.28 | 1.70 | 1.00 | 0.22 |
| YNSL | 520,343,174 | 51.52 | 32.37 | 16.02 | 2.00 | 0.91 | 0.22 |
| QH_H1 | 516,854,776 | 52.19 | 36.04 | 13.41 | 1.56 | 0.96 | 0.22 |
| QH_H2 | 507,420,565 | 51.64 | 35.44 | 13.17 | 1.84 | 0.97 | 0.21 |
| YH_H1 | 515,175,365 | 50.95 | 34.91 | 13.12 | 1.68 | 1.02 | 0.23 |
| YH_H2 | 507,527,487 | 52.17 | 35.91 | 13.55 | 1.52 | 0.97 | 0.22 |

**Supplementary Table 4. The 24 pear genomes for pan-genome analyses**

| Genome | Species | Source |
|---|---|---|
| YL | *Pyrus pyrifolia* | this study |
| ML | *Pyrus* sp. | this study |
| J20 | *Pyrus pyrifolia* | this study |
| NH | *Pyrus ussuriensis* | this study |
| HCL | *Pyrus pashia* | this study |
| BL | *Pyrus communis* | this study |
| YNSL | *Pyrus* sp. | this study |
| QH_H1 | *Pyrus pyrifolia* | this study |
| QH_H2 | *Pyrus pyrifolia* | this study |
| YH_H1 | *Pyrus pyrifolia* | this study |
| YH_H2 | *Pyrus pyrifolia* | this study |
| CG | *Pyrus pyrifolia* | https://ngdc.cncb.ac.cn/gwh/Assembly/18534/show |
| DL | *Pyrus betulifolia* | https://ngdc.cncb.ac.cn/gwh/Assembly/647/show |
| ZA1 | *Pyrus* sp. | https://doi.org/10.6084/m9.figshare.c.4502327 |
| DSS | *Pyrus pyrifolia* | https://doi.org/10.6084/m9.figshare.25139555 |
| CFLS | *Pyrus communis* | https://doi.org/10.6084/m9.figshare.25139555 |
| YLX_H1 | *Pyrus sinkiangensis* | https://pearomics.njau.edu.cn/genome_data |
| YLX_H2 | *Pyrus pyrifolia* | https://pearomics.njau.edu.cn/genome_data |
| HXS_H1 | *Pyrus sinkiangensis* | https://pearomics.njau.edu.cn/genome_data |
| HXS_H2 | *Pyrus pyrifolia* | https://pearomics.njau.edu.cn/genome_data |
| AJ_H1 | *Pyrus communis* | https://www.rosaceae.org/Analysis/17650423 |
| AJ_H2 | *Pyrus communis* | https://www.rosaceae.org/Analysis/17650423 |
| KEL | *Pyrus sinkiangensis* | https://ngdc.cncb.ac.cn/gwh/Assembly/67927/show |
| YHL1 | *Pyrus pyrifolia* | http://pyrusgdb.sdau.edu.cn/download_data.html |

**Supplementary Table 5. Primers used in this study**

| Primer | Sequence (5' to 3') |
|---|---|
| ProPpMa1-LUC-F | GGTATCGATAAGCTTATTTGCGTAAATTGCAAGTTTG |
| ProPcMa1-LUC-F | GGTATCGATAAGCTTATATACCAACAAGCAAGACACA |
| ProMa1-LUC-R | TTTGGCGTCTTCCATTTTTCGTCGGAGAAGAAGGAT |
| DAM1-SNP-Clone-F | ACGATCTCGTACAAAACCGCA |
| DAM1-SNP-Clone-R | GTCTAACAGGCCAACCCTGC |
| DAM1-SNP-Sequencing | GGATATAATTTTATTTTTATG |
| dam1-1-KASP-F1 | GAAGGTGACCAAGTTCATGCTTACCAAGGATGTGATTGCAAGGTAC |
| dam1-1-KASP-F2 | GAAGGTCGGAGTCAACGGATTTACCAAGGATGTGATTGCAAGGTAA |
| dam1-1-KASP-R | ATTTGATCCGATTTTTCCCCACCAGTATG |
| EXPA7-DNAqPCR-F | GAGGAGCTTGCGGTTATGGA |
| EXPA7-DNAqPCR-R | AGCATGCACTTGTCCCGTAA |
| NOR1-DNAqPCR-F | AAGTACCCGAACGGAGCGAGA |
| NOR1-DNAqPCR-R | GGCTTGTTGTTGGGCTTGTT |
| ProNADP-ME3-LUC-F | GGTATCGATAAGCTTGTATACCACACATACATACT |
| ProNADP-ME3-LUC-R | TTTGGCGTCTTCCATCCTCAACGACCCAAGTAATAT |
| ProARF5-LUC-F | GGTATCGATAAGCTTGAAGATATGTACTGCGACGAG |
| ProARF5-LUC-R | TTTGGCGTCTTCCATCTTCATCTCATCAAGCAAGC |
| ProAGL61-LUC-F | GGTATCGATAAGCTTCGACTGGTGGAATCATCTAAT |
| ProAGL61-LUC-R | TTTGGCGTCTTCCATGATGATTCGTATATAATCTG |
